# Drivers of Transcriptional Variance in Human Intestinal Epithelial Organoids

**DOI:** 10.1101/2021.06.02.446644

**Authors:** Zachary K. Criss, Nobel Bhasin, Sara C. Di Rienzi, Anubama Rajan, Kali Deans-Fielder, Ganesh Swaminathan, Nabiollah Kamyabi, Xi-Lei Zeng, Deepavali Chakravarti, Clarissa Estrella, Xiaomin Yu, Ketki Patil, James C. Fleet, Michael P. Verzi, Sylvia Christakos, Michael A. Helmrath, Sumimasa Arimura, Ronald A. DePinho, Robert Britton, Anthony Maresso, Jane Grande-Allen, Sarah E. Blutt, Sue E. Crawford, Mary K. Estes, Sasirekha Ramani, Noah F. Shroyer

## Abstract

**Background & Aims:** Human intestinal epithelial organoids (enteroids and colonoids) are tissue cultures used for understanding the physiology of the intestinal epithelium. Here, we explored the effect on the transcriptome of common variations in culture methods, including extracellular matrix substrate, format, tissue segment, differentiation status, and patient heterogeneity.

**Methods:** RNA-sequencing datasets from 251 experiments performed on 35 human enteroid and colonoid lines from 28 patients were aggregated from several groups in the Texas Medical Center. DESeq2 and Gene Set Enrichment Analysis (GSEA) was used to identify differentially expressed genes and enriched of pathways.

**Results:** PERMANOVA, Pearson correlations, and dendrogram analysis of all data indicated three tiers of influence of culture methods on transcriptomic variation: substrate (collagen vs. Matrigel) and format (3D, transwell, and monolayer) had the largest effect (7,271-1,305 differentially expressed genes-DEGs); segment of origin (duodenum, jejunum, ileum, colon) and differentiation status had a moderate effect (5,977-420 DEGs), and patient heterogeneity and specific experimental manipulations (e.g., pathogen infection) had the smallest effect. GSEA identified hundreds of pathways that varied between culture methods, such as IL1 cytokine signaling enriched in transwell vs. monolayer cultures, and cholesterol biosynthesis genes enriched in Matrigel vs. collagen cultures.

**Conclusions:** Surprisingly large differences in organoid transcriptome were driven by variations in culture methods such as format and substrate, whereas experimental manipulations such as infection had modest effects. These results show that common variations in culture conditions can have large effects on intestinal organoids and should be accounted for when designing experiments and comparing results between laboratories. Our data constitute the largest RNA-seq dataset interrogating human intestinal organoids.

## Introduction

The Human intestine is a highly dynamic organ that is essential for life and subject to common diseases such as cancer, infection, and chronic inflammation. The epithelial lining of the luminal surface of the intestines confers segment-specific functions to these organs: digestion and absorption of nutrients primarily occurs in the proximal small intestine (duodenum and jejunum), while the distal small intestine (ileum) absorbs remaining nutrients, bile acids, and B12; the colon/large intestine is primarily responsible for absorbing water and electrolytes and hosting the gut microbiome. The small intestinal mucosa consists of the proliferative crypt and differentiated villus compartments, whereas the epithelium of the large intestine is made up of crypts with a lower proliferative compartment and an upper differentiated compartment. Intestinal stem cells (ISCs) are located in the crypt base, where they rapidly renew and give rise to transit amplifying cells (TAC) which further differentiate into the various specialized cells of the epithelium (Barker, 2014; Barker, De Wetering, & Clevers, 2008; Scoville, Sato, He, & Li, 2008; Van Der Flier & Clevers, 2009). Paneth cells, primarily present in the small intestine, secrete antimicrobial peptides and provide signals for stem cell renewal and maintenance; goblet cells produce mucus; enteroendocrine cells release hormones; tuft cells are chemosensory in nature and also regulate immune responses; and enterocytes/colonocytes are responsible for the uptake of nutrients, water and electrolytes (Clevers & Bevins, 2013; Manley & Capecchi, 1995; Rodríguez-Colman et al., 2017; Vermeulen & Snippert, 2014). Apart from Paneth and stem cells, which remain at the crypt base, cells migrate up the crypt and differentiate prior to migrating onto the villus/colonic surface where they reside for 3-5 days before undergoing anoikis and being extruded into the lumen (Blander, 2016; Williams et al., 2015).

Experimental systems that recapitulate intestinal organs and can be cultured indefinitely are being validated and becoming vital for the progression of biomedical research. Small and large intestinal epithelial organoids (enteroids and colonoids respectively) provide researchers with an experimental platform composed of the epithelial cells and a simulacrum of the intestinal microenvironment similar to that seen *in vivo,* and thus provide excellent models for both basic and translational inquiries. Self-renewing mouse and human enteroids reported by Clevers’ lab in 2009 and 2011 replicate the polarized monolayer of the intestinal epithelium in a 3-dimensional (3D) culture (Sato et al., 2011, 2009). Human enteroids/colonoids are derived by isolating crypts from endoscopic biopsy or surgically resected tissue. Purified crypts are collected and embedded in a 3D matrix and cultured in a media containing key growth factors (Epidermal growth factor (EGF), WNT3A, R-SPONDIN, and NOGGIN) for intestinal stem cells (ISCs) renewal and maintenance (Barker, 2014; Leushacke & Barker, 2014; Middendorp et al., 2014; Sato & Clevers, 2013; Sato et al., 2009) (Figure 1A).

**Figure 1.**
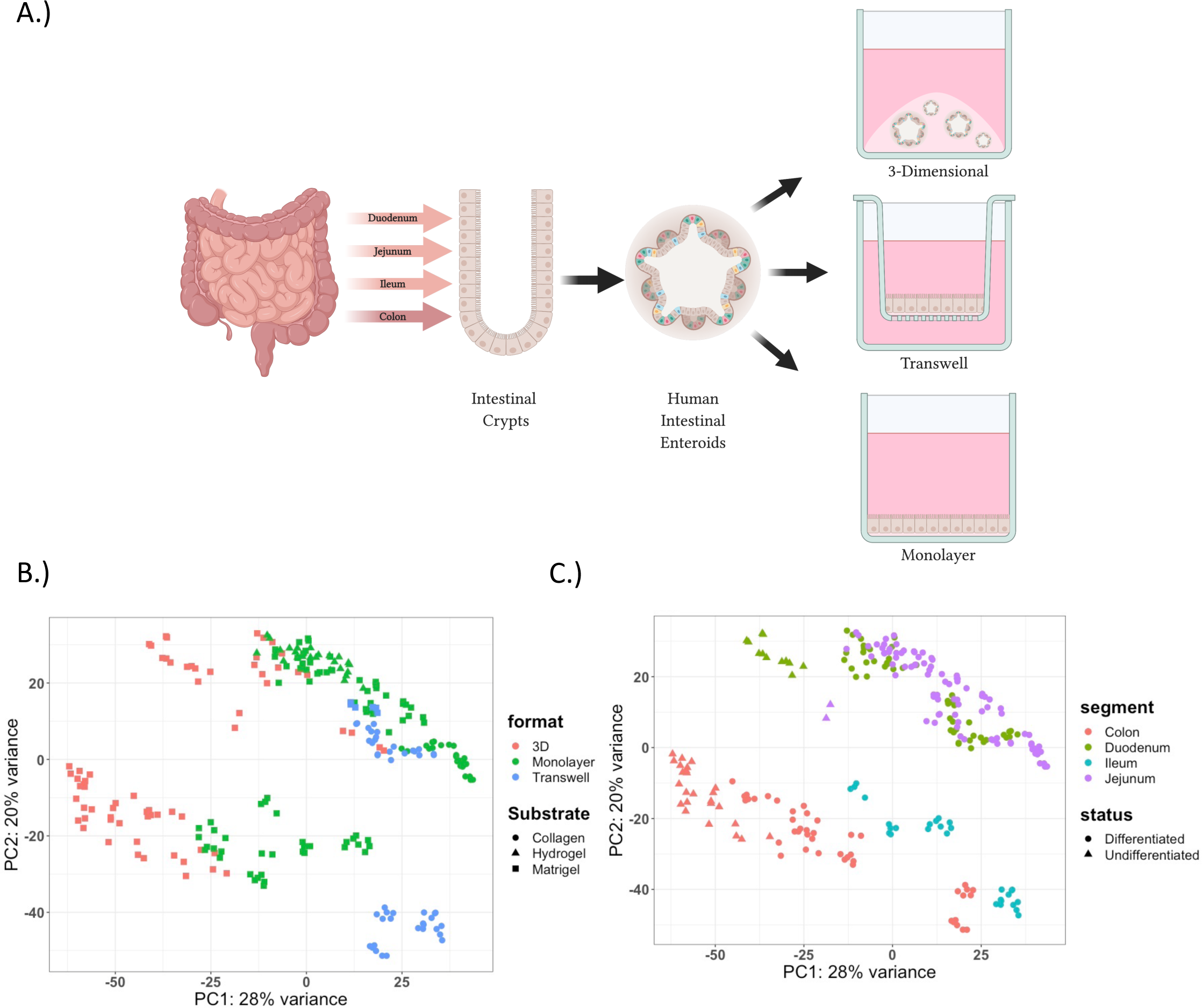
Generation of enteroid models and RNA-Seq data. A.) Schematic overview showing that 3D enteroids and colonoids are derived from crypts from various segments of the intestine. These 3D enteroids and colonoids are then able to be maintained in their 3D conformation or plated and grown in 2D as transwells or monolayers; B.) Principal component analysis (PCA) of all RNA-sequencing samples included in the analyzed dataset (n=251) labeled by the format of enteroids/colonoids (color) and the substrate used (shape): 3D-Matrigel (n=69), Monolayer-Collagen (n=24), Monolayer-Hydrogel (n=26), Monolayer-Matrigel (n=80), Transwell-Collagen (n=46) and Transwell-Matrigel (n=6); C.) PCA of all RNA-sequencing samples included in the analyzed dataset (n=251) labeled by segment of derivation (color) and differentiation status (shape): Duodenum-Differentiated (n=49), Duodenum-Undifferentiated (n=12), Jejunum-Differentiated (n=90), Jejunum-Undifferentiated (n=2), Ileum-Differentiated (n=29), Colon-Differentiated (n=44) and Colon-Undifferentiated (n=25); format: Growth formats (3D, Monolayers and Transwells), Substrate (Matrigel, Collagen IV and Matrigel-Coated Hydrogels), segment: segment organoids originated from (Duodenum, Jejunum, Ileum and Colon), status: differentiation status (Differentiated and Undifferentiated)(n=251).

This organoid model closely resembles the morphology, functionality, and cellular composition of the intestine *in vivo*, and recapitulates the expression of segment specific markers of the tissue from which they are derived (Cramer, Thompson, Geskin, Laframboise, & Lagasse, 2015; Middendorp et al., 2014; Phipson et al., 2019; Sato et al., 2009). These properties contribute another unique aspect of this model allowing for the exploration of segment specific functions or responses *ex vivo*. Enteroids/colonoids can also be grown in different “formats”, either three-dimensionally (3D) or two-dimensionally (2D), dependent on the requirements for experimentation and need for access to the apical or basal surfaces (Figure 1A). 3D enteroid/colonoid cultures are generally grown embedded in an extracellular matrix (ECM) substitute, most commonly Matrigel. These cultures are generally oriented with the apical surface of the cells facing inward and are anchored by basolateral attachments to the ECM. While 3D cultures are widely used, limited access to the luminal surface present challenges for studies such as nutrient absorption and pathogen interactions, except in the case of no ECM which results in an apical-out enteroid (Co et al., 2019; Ettayebi et al., 2016; Kozuka et al., 2017; Liu, Qi, Li, Du, & Chen, 2018; Puzan, Hosic, Ghio, & Koppes, 2018; Rajan et al., 2020; Scott et al., 2016; Thorne et al., 2018; Tong et al., 2018). 2D cultures can be grown as monolayers on ECM or transwells on ECM coated inserts (Jabaji et al., 2014; Kim, Spence, & Takayama, 2017; Liu et al., 2018; Sato et al., 2009; Scott et al., 2016; Thorne et al., 2018; Tong et al., 2018). The epithelial layer is polarized with the basolateral surface attaching to the underlying ECM (Matrigel, collagen or hydrogels) and the apical surface directed up. This allows for the apical surface to be easily accessed for experimental manipulations. The ability to change the format of enteroids/colonoids allows for the opportunity to study the influence of stimuli of the apical and basolateral surfaces with a subsequent response of the human intestinal epithelium.

The maintenance and expansion of enteroids is achieved using proliferation media conditions that promote the growth of ISCs and suppress the differentiation of the stem cells, recapitulating the intestinal crypt compartment. In addition, enteroids can be differentiated by the withdrawal of WNT3A and R-SPONDIN, and reduced NOGGIN (key factors in maintaining the stem cell state) resulting in the terminal differentiation of ISCs into absorptive and secretory cell types recapitulating the upper-crypt or villus compartment. Typically, samples grown as monolayers or on transwells are differentiated before experimental manipulation in order to mimic the in vivo interaction between the upper-differentiated compartment of the intestinal epithelium and microbes, pathogens or metabolites. Conversely, the undifferentiated crypt compartment is recapitulated through the continued growth of enteroids in proliferation media conditions. These two conditions also provide the opportunity to not only study the baseline differences between the differentiated and undifferentiated compartments but also to understand the differences in their response to particular stimuli and provide insight into compartmental specific responses and pathway enrichment.

The enteroid/colonoid model is a reproducible, scalable and physiologically relevant system applicable for a variety of high throughput screens and novel applications similar to the previous conventional culture of immortalized or cancer cell lines which lack physiological relevance (Dekkers et al., 2016, 2013; Sato & Clevers, 2013; Wilding & Bodmer, 2014). Though the enteroid/colonoid model is a significant step forward compared to the conventional cancer or immortalized cell lines, there exist differences in methods of culture that have yet to be fully compared and characterized. Here, we aggregated 251 RNA-Sequencing datasets and investigated the effect of format, substrate, segment of origin, differentiation status and patient line on driving transcriptional variance in human enteroids/colonoids.

## Results

### Format and substrate contribute the most variation to transcriptomic data from 251 human intestinal epithelial organoids samples

In order to analyze the drivers of transcriptomic variation in enteroids/organoids, we aggregated RNASeq data from 251 human enteroid/colonoid samples. These enteroids/colonoids were propagated in 3D growth conditions and replated into 3D, Transwell or Monolayer formats (Figure 1A). Subsequently, these were differentiated (where indicated) and subjected to a variety of experimental manipulations including but not limited to exposure to calcitriol (the active form of Vitamin D), human rotavirus, *E. coli* and human norovirus. Organoids were then collected, RNA extracted, and RNA-sequencing was performed resulting in a total of 251 enteroid/colonoid samples that were included in this study (Table 1). Several of these datasets have previously been published or are in preparation (Chang-Graham et al., 2019; Li et al., 2020; Lin et al., 2020; Rajan et al., 2020; Saxena et al., 2017). The organoids used were derived from various segments (duodenum, jejunum, ileum or colon), from various patients (28 patients), grown in various formats (3D, monolayers on 96-well plates (monolayers) or monolayers on transwells (transwells)) and substrates (Collagen IV, Matrigel or Matrigel-coated Hydrogels), grown under different differentiation statuses (differentiated and undifferentiated) and given different experimental treatments (20), which were performed by different experimenters/projects (10) (Table S1).

**Table 1.** 251 RNA-Sequencing Samples used in this analysis. Shown are the abbreviated demographics with the number of Formats (3D, Monolayers and Transwells), Substrates (Matrigel, Collagen IV and Matrigel-Coated Hydrogels), Statuses (Differentiated and Undifferentiated), Lines and Treatments for enteroid/colonoid lines derived from every segment (n=251).

|  | Duodenum | Jejunum | Ileum | Colon |
| --- | --- | --- | --- | --- |
| Formats | 3 | 3 | 2 | 3 |
| Substrates | 2 | 3 | 2 | 2 |
| Statuses | 2 | 2 | 1 | 2 |
| Lines | 8 | 4 | 3 | 20 |
| Treatments | 13 | 15 | 10 | 12 |
| # of Samples | 61 | 92 | 29 | 69 |

To begin investigating the drivers of variance among samples, we performed Permutational multivariate analysis of variance (PERMANOVA) on our nonparametric dataset (Table 2). These revealed that the top two drivers of variance in this large dataset were format and substrate (F = 90.6 and 80.4, respectively), which accounted for 21.3% and 18.9%, respectively, of the variation in our model. Segment and differentiation status were moderate drivers of variance (F = 23.2 and 19.8, respectively) and explained 8.2% and 2.3%, respectively, of the model, and finally line (representing patient-to-patient variation), and experimental treatment (including various experimental treatments such as infection and calcitriol treatment; see methods for details) had the smallest contribution to the variance (F = 4.9 and 4.5, respectively) and explained 15.5% and 10.6% of the variance in the model respectively).

**Table 2.**
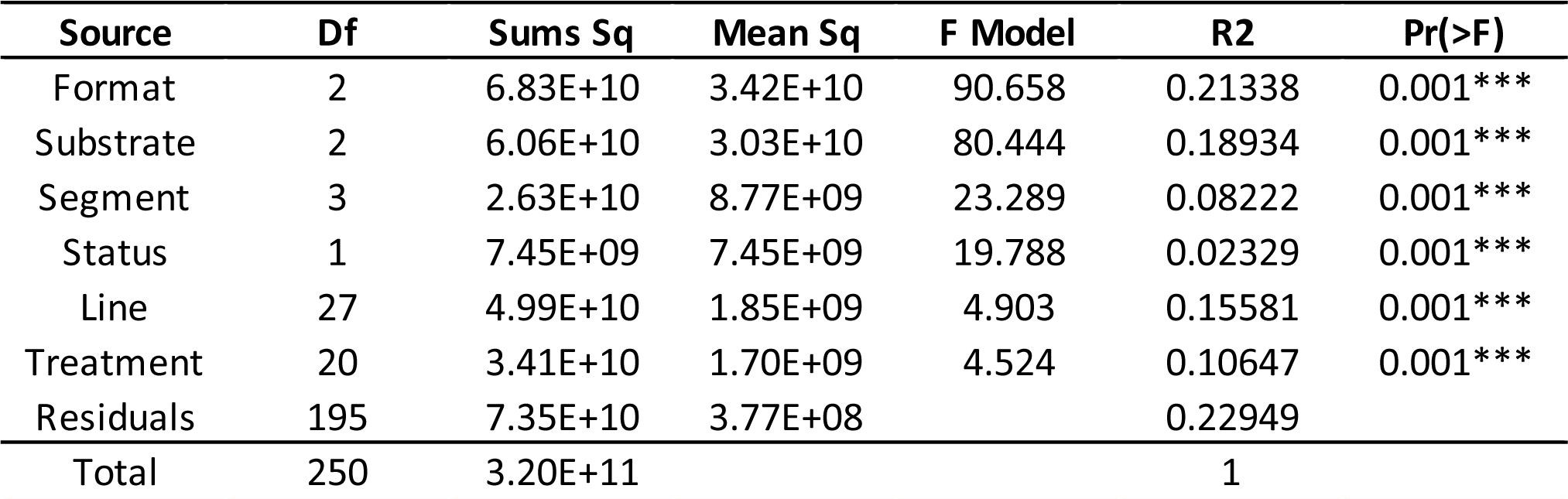
PERMANOVA Results. PERMANOVA output showing the effects of Format, Substrate, Segment (segmental origin of enteroid/colonoid), Status (differentiation status), Line (patient line), and Treatment based on Euclidian distances and 999 permutations. Df: degrees of freedom; Sums Sq: sum of squares; Mean Sq: mean sum of squares; F Model: *F* statistics; R2: partial *R-squared;* Pr (>F): *P* values, *** = 0.001;

| Source | Df | Sums Sq | Mean Sq | F Model | R2 | Pr(>F) |
| --- | --- | --- | --- | --- | --- | --- |
| Format | 2 | 6.83E+10 | 3.42E+10 | 90.658 | 0.21338 | 0.001*** |
| Substrate | 2 | 6.06E+10 | 3.03E+10 | 80.444 | 0.18934 | 0.001*** |
| Segment | 3 | 2.63E+10 | 8.77E+09 | 23.289 | 0.08222 | 0.001*** |
| Status | 1 | 7.45E+09 | 7.45E+09 | 19.788 | 0.02329 | 0.001*** |
| Line | 27 | 4.99E+10 | 1.85E+09 | 4.903 | 0.15581 | 0.001*** |
| Treatment | 20 | 3.41E+10 | 1.70E+09 | 4.524 | 0.10647 | 0.001*** |
| Residuals | 195 | 7.35E+10 | 3.77E+08 |  | 0.22949 |  |
| Total | 250 | 3.20E+11 |  |  | 1 |  |

We next visualized the effect of these sources of variation of the data using Principal Component Analysis (PCA) (Figure 1B & 1C). Principal Component Analysis (PCA) of the entire dataset was used to visualize clustering of the samples, with the first two principal components accounting for 29% and 19% variance of the samples. It was observed that enteroids/colonoids grown in 3D (red) were left-shifted on the y-axis compared to samples from the same segment grown in 2D (transwells (blue) and monolayers (green)) (Figure 1B). The PCA plot exhibited that duodenal (green) and jejunal (purple) samples clustered together in the first and second quadrant while the colon (red) and ileal (blue) samples existed in the third and fourth quadrant respectively. With undifferentiated enteroids/colonoids being left-shifted on the y-axis compared to differentiated samples (Figure 1C).

### 2D human enteroid substrates are a major driver of variation

In order to further examine the drivers of transcriptomic variance, we investigated the variance across the dataset subset by segment of origin (duodenum, jejunum, ileum and colon). Using PCA analysis of datasets derived from individual segments, we observed that the samples clustered by format and substrate (Figure 2A, S1, S2 & S3). For example, in duodenum samples there are 4 observable clusters: monolayers grown on Matrigel, transwells and monolayers grown on collagen, and two 3D clusters grown in Matrigel (Figure 2A). PC1 in this analysis, accounting for 38% variance, appears to be driven by the dimensionality of the format, with the 2D enteroids clustered in the first and fourth quadrants of the plot while the 3D enteroids clustered in the second and third quadrant of the PCA plot (Figure 2A). PC2, accounting for 20% variance, appears to be driven primarily by substrate with monolayers grown on Matrigel clustering in the first quadrant of the plot and transwells and monolayers grown on collagen clustering in the fourth quadrant of the PCA plot. Furthermore, sub-clusters within the fourth quadrant separate monolayer and transwell samples grown on collagen. The separation of 3D samples into two clusters is driven by their differentiation status, examined in greater detail below. Additionally, a dendrogram of the data shows a clear bifurcation of the duodenal monolayers based on the substrate regardless of the line or other experimental manipulations (Figure 2B). This same observation was observed in a Pearson correlation matrix of duodenal enteroids (Figure S9). Patient-to-patient variation and the experimental treatments (e.g., infection) had small effects on the variation within the dataset, which could be observed in the dendrograms as the final branch points in the tree, all reinforcing the results of the PERMANOVA analysis of our data (Figure 2B and Table 2). Similar observations were made by analyzing clustering of jejunal, ileal, and colonic datasets using PCA, dendrogram and Pearson correlation analysis which demonstrated clustering of samples by format and substrate (Figures S1A-B, S2A-B, S3A-B and S9-12). Additionally, in the jejunum, monolayer samples were also grown on synthetic hydrogel bases that were coated with Matrigel (thus providing different stiffness to the Matrigel substrate). Data from these monolayers that were grown on hydrogels clustered together with other monolayer samples grown on Matrigel-coated plastic, with distinct subclusters associated with differing hydrogel properties (Figure S1A-B). In-depth analysis of the influence of substrate stiffness on enteroid physiology will be published by Grande-Allen et al. (manuscript in revision); however, these results underscore that the composition of the substrate matrix is a major driver of variance.

**Figure 2.**
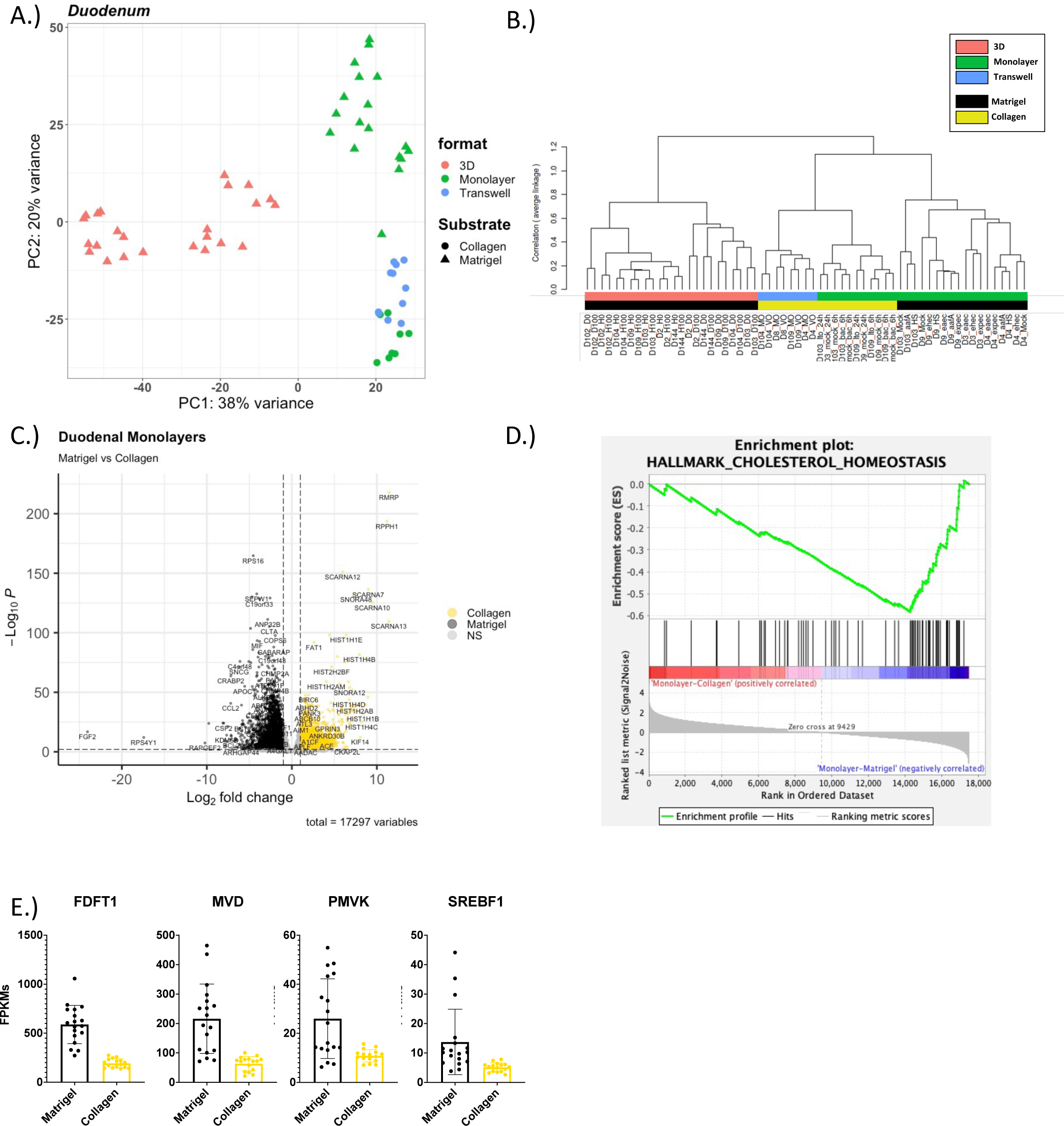
Cholesterol biosynthesis related genes are upregulated in duodenal monolayers grown on Matrigel. A.) PCA of the RNA-sequencing datasets for duodenal enteroids: 3D-Matrigel (n=24), Monolayer-Matrigel (n=18), Monolayer-Collagen (n=8), and Transwell-Collagen (n=11); B.) A dendrogram with agglomerative hierarchal clustering of the duodenal gene set from RNA-sequencing. Branch length indicates degree of difference between samples; C.) Volcano plot of differentially expressed genes when comparing duodenal monolayers on collagen (gold) and Matrigel (black). Gold/black dots indicate differentially expressed genes (FDR ≤ 0.01 and a foldchange ≥2 or ≤0.5) that are upregulated or downregulated (respectively) in duodenal monolayers on collagen (3136 genes) vs. duodenal monolayers on Matrigel (2868 genes); D.) Gene Set Enrichment Analysis (GSEA) showing an enrichment of the hallmark cholesterol homeostasis get set signature in duodenal monolayers grown on Matrigel when compared to duodenal monolayers grown on collagen; E.) Normalized Fragments Per Kilobase of transcript per Million mapped reads (FPKMs) of cholesterol biosynthesis genes from RNA-Sequencing of duodenal monolayers grown on Matrigel (black) and collagen IV (gold)s;

We used DESeq2 to identify differentially expressed genes when comparing datasets from organoids in which only one of the four major variables (segment, substrate, format, differentiation status) was different (Love, Huber, & Anders, 2014). Thus, a total of 30 comparisons were made (Table S2). Genes were considered differentially expressed with an FDR ≤ 0.01 and a fold change ≥2 or ≤0.5. We identified genes differentially regulated in response to substrate. Specifically, as visualized in a volcano plot 5244 genes were differentially expressed in duodenal enteroids grown as monolayers on collagen (3136 genes upregulated) versus Matrigel (2868 genes upregulated) (Figure 2C and Table S2). We next assessed pathways that were differentially regulated by substrate using Gene Set Enrichment Analysis (GSEA) with FDR ≤ 0.05 (Mootha et al., 2003; Subramanian et al., 2005). GSEA revealed a total of 36 of the Hallmark collection gene sets that were enriched in duodenal monolayers grown on collagen (6 gene sets) and duodenal monolayers grown on Matrigel (30 gene sets). There were also 238 GSEA canonical pathway gene sets that were enriched in duodenal monolayers grown on collagen (4 gene sets) and duodenal monolayers grown on Matrigel (234 gene sets). And 326 Gene Ontology (GO) gene sets that were enriched in duodenal monolayers grown on collagen (10 gene sets) and duodenal monolayers grown on Matrigel (326 gene sets). Similar analyses comparing jejunal monolayers grown on collagen and Matrigel identified 24 Hallmark pathway gene sets, 193 GSEA canonical pathway gene sets and 278 GO term gene sets that were enriched (FDR ≤ 0.05). Datasets from ileum and colon did not include organoids grown in the same format with different substrates, therefore a focused analysis of pathways that were driven by substrate alone is confounded by differences in format (Figure S2 and S3, Table S2).

Of interest, the cholesterol homeostasis Hallmark gene set from MSigDB was enriched in duodenal monolayers grown on Matrigel compared to duodenal monolayers grown on collagen (NES = -2.58 and FDR = 0) (Figure 2D), and this pathway was also observed in jejunal monolayers on Matrigel when compared to jejunal monolayers on collagen (NES = -1.84and FDR = 1E-3) (Figure S1C). Several genes that are involved in the biosynthesis of cholesterol (*PMVK, MVD,* and *FDFT1*) were upregulated in monolayers on Matrigel compared to collagen (Figure 2E). SREBF1 (Sterol Regulatory Element Binding Transcription Factor 1) a regulator of cholesterol biosynthesis, was also significantly upregulated in duodenal monolayers that were grown on Matrigel corresponding with the increase in enzymes that are involved in cholesterol biosynthesis (Figure 2E).

### Growth format is a major driver of variation in human enteroids and colonoids

In addition to observing the effects of the substrates on enteroid monolayers in PCA plots and dendrograms, we also observed differences between enteroids grown in different growth formats with the same substrate (Figure 2A&B and S1). DESeq2 analysis identified 4695 genes that were differentially expressed between duodenal enteroids grown on collagen in the monolayer (3378 genes upregulated) or transwell (1317 genes upregulated) formats (Figure 3A, Table S2). Thus, we further investigated monolayers and transwells that were both grown on collagen. GSEA (FDR ≤ 0.05) revealed a total of 31 of Hallmark pathways gene sets that were enriched in duodenal monolayers grown on collagen (2 gene sets) or in duodenal transwells grown on collagen (29 gene sets). There were also 505 GSEA canonical pathway gene sets that were enriched in duodenal monolayers grown on collagen (2 gene sets) or in duodenal transwells grown on collagen (503 gene sets). 400 GO gene sets were only enriched in duodenal transwells grown on collagen.

**Figure 3.**
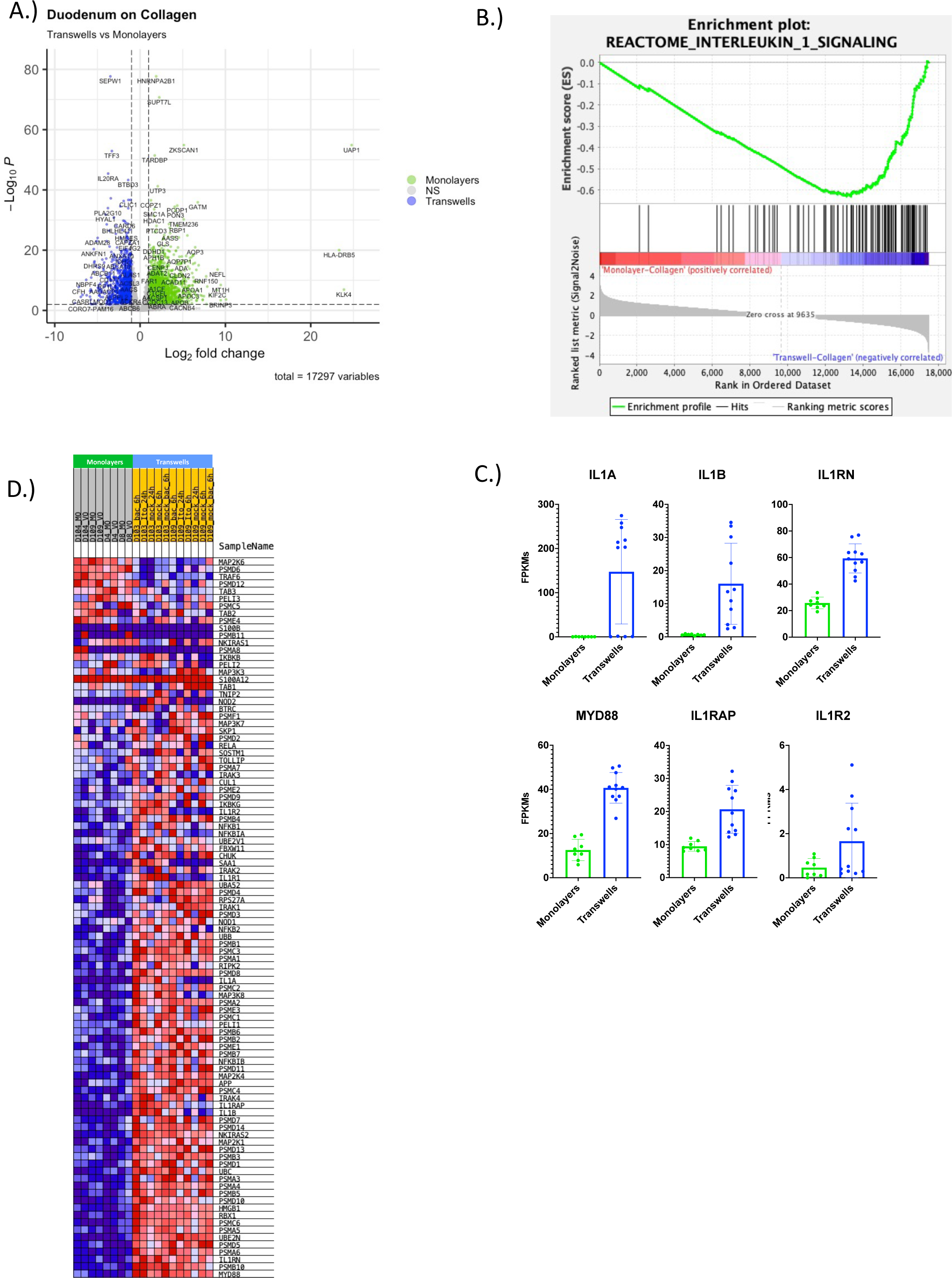
Interleukin-1 signaling pathway is enriched in duodenal transwells grown on collagen. A.) Volcano plot of differentially expressed genes when comparing duodenal monolayers (green) and transwells (blue) grown on collagen. Green/blue dots indicate differentially expressed genes (FDR ≤ 0.01 and a foldchange ≥2 or ≤0.5) that are upregulated or downregulated (respectively) in duodenal monolayers on collagen (3378 genes) vs. duodenal transwells on collagen (1317 genes); B.) GSEA showing an enrichment of the reactome interleukin-1 signaling get set signature in duodenal transwells grown on collagen when compared to duodenal monolayers grown on collagen; C.) Normalized FPKMs of Interleukin-1 signaling pathway genes that encode for secreted cytokines (*IL1A*&*IL1B*) and inhibitors (*IL1RN*) (top), and Interleukin-1 Receptor Machinery (bottom) on duodenal monolayers (green) and transwells (blue) grown on collagen; D.) GSEA showing an enrichment of the reactome interleukin-1 signaling get set signature in duodenal transwells grown on collagen when compared to duodenal monolayers grown on collagen;

Of note, the Reactome Interleukin 1 signaling gene set was enriched in duodenal transwells grown on collagen when compared to monolayers grown on collagen (NES = -3.05 and FDR = 0) (Figure 3B). Similarly, the Reactome interleukin 1 signaling gene set was also enriched in jejunal transwells grown on collagen when compared to monolayers grown on collagen (NES = -2.56 and FDR = 2.32E-04) (Figure S1D). Several of the enriched genes were associated with the receptor machinery for IL-1 signaling (*ILRAP*, *IL1R2* and *MYD88*). Moreover, secreted protein transcripts were also enriched in transwells, those being proinflammatory IL-1 cytokines (*IL1A* & *IL1B*) as well as an IL-1 receptor antagonist (*IL1RN*) (Figure 3C). We also observed an enrichment in the hypoxia hallmark gene set in duodenal (NES = -1.72 and FDR = 0.001) and jejunal (NES = 1.78 and FDR = 0.003) transwells grown on collagen compared to monolayers grown on collagen (Figure S5).

Additionally, a comparison of 3D and monolayers grown on/in Matrigel revealed several genes and gene sets that were differentially expressed. These comparisons were made in duodenal, jejunal and colonic organoids (Table 2A & Figure S4). DESeq2 analysis identified genes that were differentially expressed between duodenal (5139 genes), jejunal (5195 genes) and colonic (6049 genes) organoids grown in 3D (duodenum-2615, jejunum-1569 and colon-3481 upregulated genes) or on monolayers (duodenum-2524, jejunum-3635 and colon-2568 upregulated genes) with Matrigel (Table 2A). Additionally, the Hallmark WNT Beta Catenin signaling gene set was enriched for 3D organoids in Matrigel compared to monolayers on Matrigel (Figure S4A). Additionally, the GO Transmembrane receptor protein kinase activity gene set was also enriched for 3D organoids grown in Matrigel compared to monolayers on Matrigel (Figure S4B). Several of the GO Transmembrane receptor protein kinase activity genes that were upregulated in 3D enteroids were related to the proliferative and crypt compartment such as EphB2, EphB3 (Figure S4C) (Merlos-Suárez et al., 2011). We also observed an enrichment for the Reactome antigen processing and cross presentation gene set in colonoid monolayers grown on Matrigel compared to 3D colonoids in Matrigel (Figure S3C-D). Colonoid monolayers on Matrigel had an enrichment specifically in the HLA-E transcripts, a nonclassical MHC I molecule (Perera et al., 2007; Shao, Kamalu, & Mayer, 2005). In another case we observed an enrichment of several ABC transporters, known for their role in the transport of molecules and drug resistance, in ileal transwells on collagen compared to ileal monolayers on Matrigel (Figure S2C-D) (Mutch et al., 2004; Tsai et al., 2017).

### Known transcriptional regional identity markers are observed in enteroids

The human intestine is comprised of several specialized segments with dedicated functional capabilities. Gene expression profiles of each of the segments has been characterized in previous reports (Camp et al., 2014; Yu, Mu, Yang, Su, & Zhu, 2017; Zheng et al., 2015). To highlight the variation in gene expression between human enteroids derived from different segments of the intestine (duodenum, jejunum, ileum and colon), we performed PCA of enteroids/colonoids that were grown using the same format (3D, monolayer, or transwell). PCA of organoids cultured on transwells only showed that colonic and ileal samples were separated from the jejunal and duodenal samples by PC1 on the x-axis, accounting for 50% variance in this dataset. Specifically, the ileal and colonic samples clustered in the second and third quadrant, respectively, on the PCA plot. The duodenal and jejunal samples clustered in the first and fourth quadrant of the PCA plot, suggesting a higher level of similarity between duodenal and jejunal samples than ileal and colon samples (Figure 4A). This observation was replicated when using a dendrogram which showed clear separations between the colonic and ileal samples being broadly separated from the duodenal and jejunal samples that were intermingled (Figure 4B). Similar observations could also be made across formats, where samples derived from duodenum and jejunum clustered together and samples from ileum and colon also clustered together (Figure 4A-B & S6A-D). As previously mentioned, it was observed that the substrate that the enteroids were grown on is a large driver of variance, in this case appears to be the driver of PC2 and a delineator of jejunal and duodenal samples in the dendrogram (Figure 4A-B). Additionally, the jejunal enteroids that were grown on Matrigel (ULDM1 & ULDM2) were also transduced with a doxycycline-inducible neurogenin-3 (*NGN3*) (uninduced) construct, an additional factor that may have driven the differences between other jejunal samples.

**Figure 4.**
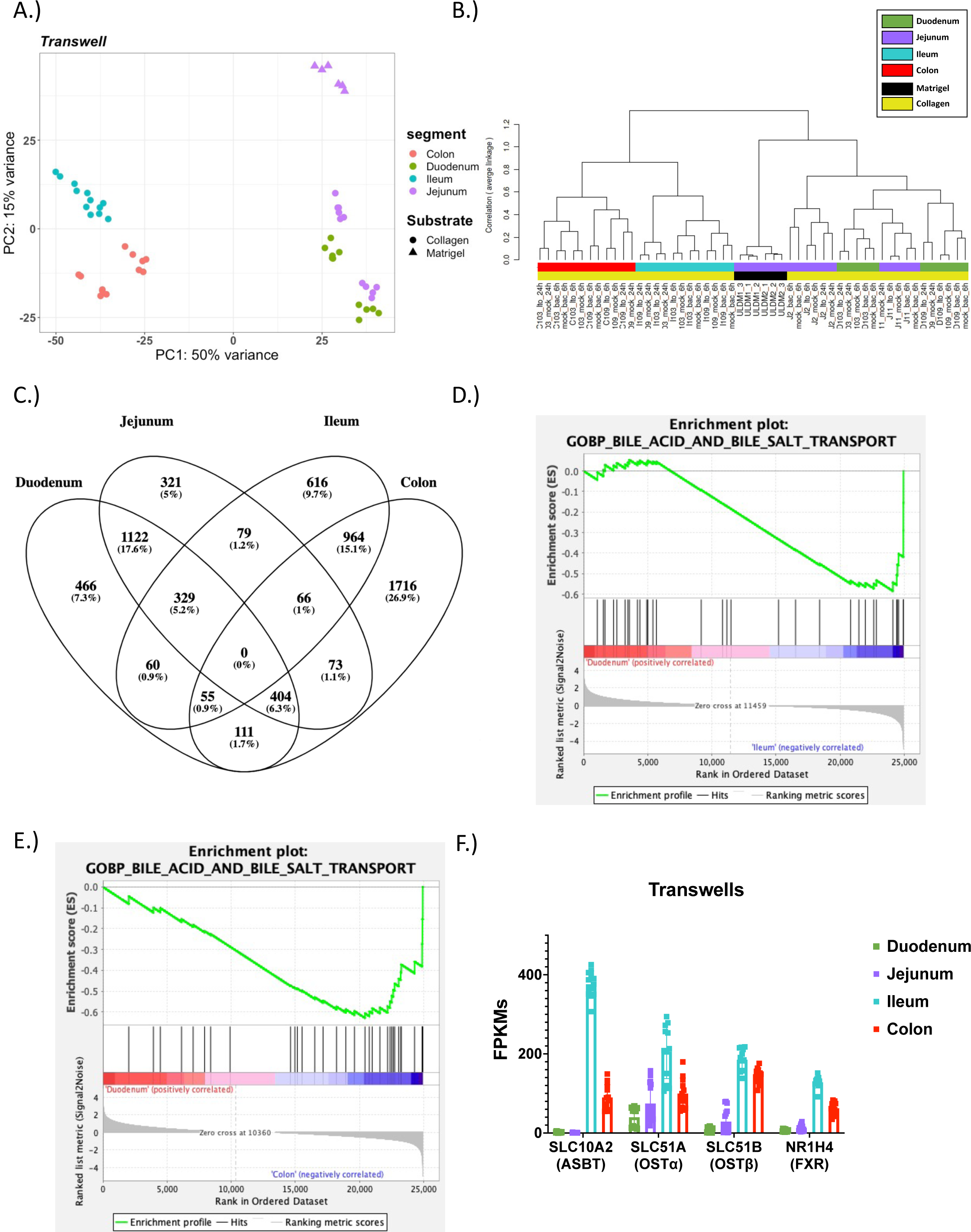
Bile acid transport machinery is enriched in ileal enteroids and colonoids. A.) PCA of the RNA-sequencing datasets for enteroids on transwells: Duodenum-Collagen (n=11), Jejunum-Collagen (n=11), Jejunum-Matrigel (n=6), Ileum-Collagen (n=12), and Colon-Collagen (n=12); B.) A dendrogram with agglomerative hierarchal clustering of the transwell gene set from RNA-sequencing. Branch length indicates degree of difference between samples; C.) Venn diagram displaying the overlap of differentially expressed genes (DEGs) between duodenal, jejunal, ileal and colonic intestinal organoids grown on transwells. DEGs were were defined as any gene that was differentially enriched in the segment of interest compared to any other segment with DESeq2 (FDR ≤ 0.01 & Fold Change ≥2 and ≤ 0.5); D & E.) GSEA showing an enrichment of the Gene Ontology (GO) bile acid and bile salt transport get set signature in (D) ileal enteroids and (E) colonoids on transwells when compared to duodenal enteroids grown on transwells; F.) Normalized FPKMs of bile acid transport genes in duodenal (green), jejunal (purple), ileal (blue) and colonic (red) intestinal organoids grown on transwells;

We utilized DESeq2 analysis to generate differentially expressed gene lists between segments within the same format. Samples were considered differentially expressed with an FDR ≤ 0.01 & fold changes ≥2 or ≤0.5 (Table S2). In order to show the similarities and differences of intestinal organoids derived from various segments we generated Venn diagrams using the DEGs for each segment (Figure 4C and S6E). The DEGs list for each segment was composed of all enriched genes in the segment of interest compared to any other segment (e.g., union list of all duodenal enriched genes in Duodenum vs. Jejunum, Duodenum vs. Ileum, and Duodenum vs. Colon). As indicated by the PCA plot the duodenum and jejunum (1,122 shared DE genes) shared the most similarity and the ileum and colon (964 shared DE genes) were also similar to each other (Figure 4C).

To understand the potential phenotypic impact due to the differences in gene expression between segments, GSEA was performed (FDR ≤ 0.05) (Figure 4D-F, Supplementary Information 3). GSEA revealed a total of 92 of GO gene sets that were enriched in duodenal transwells grown on collagen (14 gene sets) and ileum transwells grown on collagen (78 gene sets). Additionally, a total of 141 GO gene sets were enriched in duodenal (133 gene sets) and colon transwells grown on collagen (8 gene sets). There was an enrichment in the bile acid and bile salt transport gene ontology term in ileal (NES = -1.61 and FDR = 0.11) and colonic (NES = -1.88 and FDR = 0.04) transwells relative to the duodenal transwells (Figure 4D-E). As anticipated, these transcripts for bile acid transport genes were upregulated in the ileum and colon compared to the duodenum and jejunum (Figure 4F).

### Differentiation status drives known changes in transcriptional markers proliferating and differentiated enteroids and colonoids

The transcriptional and functional differences between the undifferentiated crypt compartment and the upper villus/differentiated compartment of the intestine have been well characterized *in vivo* and *in vitro*. Here, we confirmed and expanded on the importance of differentiation on the enteroid/colonoid model.

As shown above, the format and substrate are major drivers of variation in enteroids, however further sub-clusters can be observed that are distinguished by differentiation status (Fig. 1C). By analyzing duodenal and colonic organoids grown in 3D, differences between the differentiated (red) and undifferentiated (cyan) cultures can be observed in PCA plots and dendrograms (Fig. 5A&B & S7A&B). The dendrogram and the first principal component of the PCA revealed a clear separation between differentiated and undifferentiated enteroids (Fig. 5A&B & S7A&B). DESeq2 analysis identified 2902 genes that were differentially expressed between duodenal enteroids that were differentiated (1474 genes enriched) and undifferentiated (1428 genes enriched). In this analysis only control samples were used due to the presence of calcitriol responsive genes that were compartment specific, which will be explored in a manuscript in preparation (Fig. 5C & Table S2). Similarly, there were 2756 genes that were differentially expressed between colonoids that were differentiated (1338 genes enriched) and undifferentiated (1418 genes enriched) (FDR ≤ 0.01 & fold changes ≥2 or ≤0.5) (Fig. S7C & Table S2). Consistent with their proliferative and self-renewing nature, markers of proliferation and intestinal stem cells were upregulated in undifferentiated enteroids and colonoids (*ASCL2*, *KI67* and *LGR5*) (Fig. 5C-D & S7C-D). Conversely, differentiated enteroids and colonoids expressed genes that are associated with differentiated cells in the duodenum (MUC2 & TFF3) and Colon (CA2 & TFF3) (Fig. 5D & S7D). Here, we explored the well-known effect of changing media conditions resulting in the shift from crypt like stem cells to villus-like differentiated cell types.

**Figure 5.**
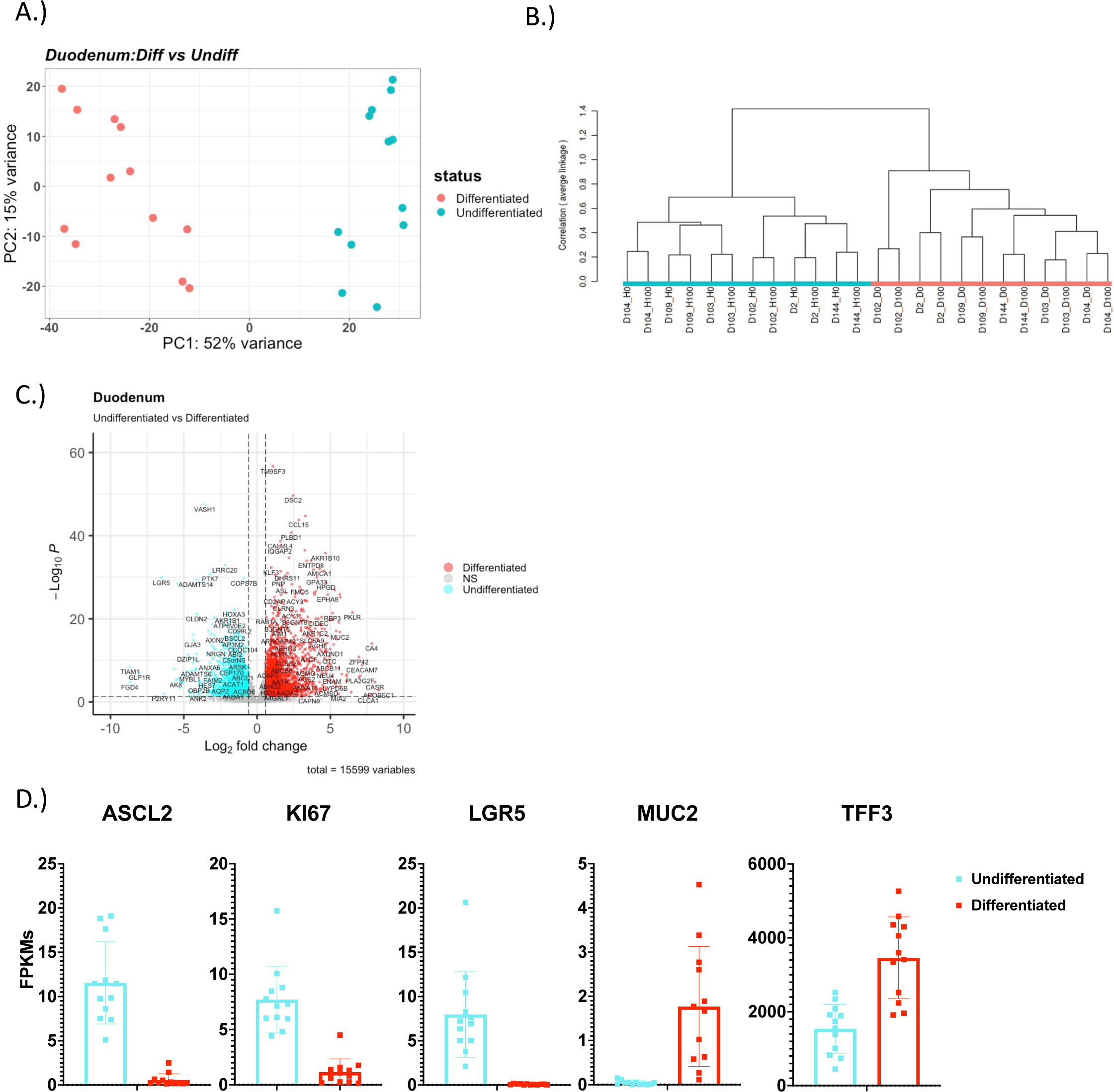
Differentiation media conditions drive changes in proliferation and differentiation markers in 3D duodenal enteroids. A.) PCA of the RNA-sequencing datasets for 3D Duodenal enteroids: Differentiated (n=12) and Undifferentiated (n=12); B.) A dendrogram with agglomerative hierarchal clustering of the 3D duodenal gene set from RNA-sequencing. Branch length indicates degree of difference between samples; C.) Volcano plot of differentially expressed genes when comparing differentiated (red) and undifferentiated (cyan) 3D duodenal enteroids. Red/cyan dots indicate differentially expressed genes (FDR ≤ 0.01 and a foldchange ≥2 or ≤0.5) that are upregulated or downregulated (respectively) in differentiated (1474 genes) vs. undifferentiated (1428 genes) 3D duodenal enteroids; D.) Normalized FPKMs of undifferentiated (cyan) and differentiated (red) 3D duodenal enteroids for stem cell (*ASCL2*, *KI67* and *LGR5*) and differentiation (*MUC2* and *TFF3*) markers;

### Patient to patient variability is a driver of variation between samples

Another variable of interest is the difference between patients. We previously described patient-to-patient variability in jejunal enteroid lines suggesting significant baseline variability between enteroids derived from different individuals, but with little variation between technical replicates (Lin et al., 2020; Saxena et al., 2017). Here, we examined patient-to-patient variation on a larger scale with a study encompassing 28 patient lines from the duodenum, jejunum, ileum and colon (Figure S1A). Analysis of differentiated 3D duodenal enteroids shows that samples from the same patient pair together (control and treatment) on a PCA plot, and the experimental treatment (in this case, exposure to calcitriol for 24 hours) results in small shifts on the X-axis on the PCA plot compared to the controls (Figure 6A). Additionally, the dendrogram shows clustering of patient lines regardless of the experimental manipulation (Figure 6B). A parallel analysis of undifferentiated 3D duodenal enteroids and colonoids shows that patient samples clustered together despite experimental manipulations (Figure 5B & S7B). Clustering by patient is also observed in samples differentiated on transwells, where samples from the same patient line cluster together as well despite undergoing several distinct experimental manipulations such as bacteria, rotavirus, calcitriol, or vehicle exposures (Figure 4A-B).

**Figure 6.**
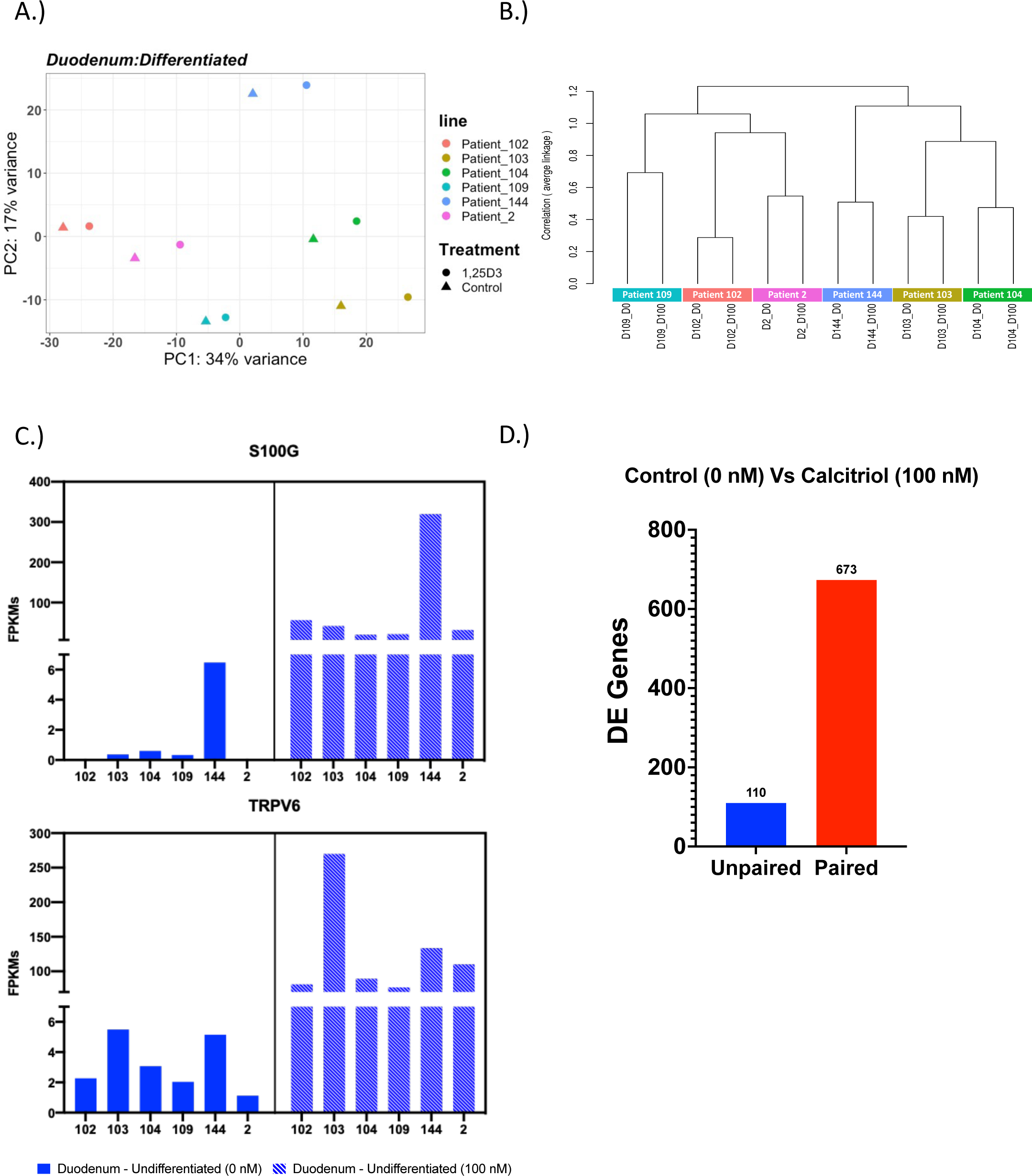
Patient-to-patient variability results in variable basal gene expression and response to stimuli. A.) PCA of the RNA-sequencing datasets for 3D Differentiated duodenal enteroids (n=12) with 6 patient samples indicated by 6 different colors; B.) A dendrogram with agglomerative hierarchal clustering of the 3D differentiated duodenal gene set from RNA-sequencing. Branch length indicates degree of difference between samples; C.) Normalized FPKMs of *S100G* (top) and *TRPV6* (bottom) expression in individual differentiated 3D duodenal enteroids lines comparing the gene expression level between the control group (0nM, solid) and the calcitriol treatment group (100nM, dashed); D.) The number of differentially expressed (DE) genes that were output from DESeq2 as a result of using an unpaired (blue) or paired (red) experimental design;

Upon further investigation of differentiated 3D duodenal enteroids we observed that basal gene expression as well as transcriptional response to calcitriol treatment were highly variable among 6 distinct patient lines (D102, D103, D104, D109, D144 and D2). Basal gene expression of *S100G,* which encodes the calcium binding protein calbindin-D9k, was highly variable with FPKM values of 0 (D102), 0.37 (D103), 0.59 (D104), 0.33 (D109), 6.47 (D144) and 0 (D2). Upon Calcitriol treatment *S100G* expression was increased proportionally with FPKM values of 56.44 (D102), 41.52 (D103), 20.01 (D104), 21.47 (D109), 319.58 (D144), and 32.02 (D2) (Figure 6C). Likewise, *TRPV6,* encoding the transporter which mediates uptake of the Ca^2+^ from the intestinal lumen to the cytosol, also showed variable expression that increased proportionally upon calcitriol treatment: FPKM values in differentiated enteroids were 2.27 (D102), 5.49 (D103), 3.06 (D104), 2.03 (D109), 5.14 (D144) and 1.12 (D2); and increased upon calcitriol treatment to 81.15 (D102), 269.82 (D103), 89.22 (D104), 76.78 (D109), 133.54 (D144) and 110.14 (D2). This same pattern of patient-specific response to treatment was also observed in undifferentiated duodenal enteroids as well as in differentiated and undifferentiated colonoids (Figure S8A-C). In-depth analysis of the influence of calcitriol on human enteroids and colonoids is in preparation (Criss et al.), however these results briefly underscore one of the points of patient-to-patient variability that is observed in different enteroid/colonoid lines.

These data demonstrate what was observed with PERMANOVA that although patient-to-patient variation in enteroids and colonoids is low compared to the influence of other variables--such as substrate, format, segment, and differentiation status—it is nevertheless significant and often greater than the experimental manipulation such as infection or hormone treatment. Because the experimental treatment of interest frequently does not drive transcriptional differences as strongly as patient-to-patient variation, we sought to compare analytical results using DESeq2 to observe the differences between the comparison of group means (unpaired) versus a paired statistical analysis. Thus, when examining differentially expressed genes in response to calcitriol treatment in differentiated duodenal enteroids, the number of DE genes was markedly higher using a paired analysis (673 genes) compared to an unpaired analysis (110 genes) (Figure 6D). Similarly, the number of genes differentially expressed in differentiated colonoids treated with calcitriol was markedly higher using a paired analysis (447 genes) versus an unpaired analysis (64 genes). This observation was consistent in undifferentiated duodenal enteroids (unpaired: 53 genes and paired: 210) and colonoids (unpaired: 16 genes and paired:. 106) (Figure S8D). Likewise, the number of differentially expressed genes in response to differentiation is also increased using a paired analysis, although the magnitude of the affect is proportionally smaller because of the large effect size of differentiation compared to patient-to-patient variation (Figure S7E).

## Discussion

Human enteroids and colonoids have been used increasingly over the past decade as *in vitro* organoid models to recapitulate human intestinal physiology and response to stimuli. However, a comprehensive examination of experimental parameters that drive transcriptomic changes in these organoids had not yet been performed. Here, we compiled over 250 RNA-Seq samples from several studies across of a consortium of laboratories to explore the effects of common variables on the transcriptome of this intestinal organoid platform. We showed that the experimental variables were able to be separated into three tiers based on magnitude of effect: 1.) growth format and substrate 2.) segment and differentiation status, and 3.) patient line and experimental treatment. Here, we explored specific examples of biological pathways that were impacted by differences in the growth format, substrate and segment of origin. We were also able to observe the well-known effect of manipulating differentiation status by growth factor withdrawal. Additionally, the importance of accounting for patient-to-patient variability was addressed when using lines derived from several patients. Moreover, lines from the same patient were found to be transcriptionally similar to each other even subsequent to experimental treatments, which were shown to have the smallest impact on the transcriptome. From this study a strong case is made that the substrate the enteroids and colonoids are grown on should be taken into consideration when designing experiments. Here, we observed that within the same segment and format, the substrate that enteroids are grown on is a large driver of variance in 2D and potentially 3D cultures (Figure 1D, 2, S1 & S3). A recent study performed in 3D enteroids showed that the loss of contact with ECM results in re-oriented, apical-out enteroids, underscoring the importance of the substrate in establishing orientation and cell polarity (Co et al., 2019). Among thousands of gene expression differences, we found that duodenal and jejunal monolayers grown on Matrigel both had increased expression of the cholesterol biosynthesis pathway compared to those grown on collagen. We speculate that the consequential effect of the substrate is either due to the transduction of mechanical forces to the cells or the presence of a ligand that induces transcriptional changes. *In vivo* it is known that the basal surface of the intestinal epithelium binds to and interacts with the underlying basement membrane via integrins that are differentially expressed along the crypt and villus axis (Lussier, Basora, Bouatrouss, & Beaulieu, 2000). This binding of integrins to the ECM components of the basement membrane is known to result in a mechanical stress on the epithelium (Tang, 2020). Mechanical stress on the intestine has been shown to result in transcriptional changes, for instance an *in vivo* model of cancer exhibited that a prolonged mechanical pressure resulted in the nuclear translocation of β-catenin in the intestine (Fernández-Sánchez et al., 2015). *In vitro* substrate stiffness has also been shown to affect the proliferation and migration of caco-2 cells, a colorectal adenocarcinoma cell line, as well in other cell types (Du et al., 2016; Gjorevski et al., 2016; M. A. Sanders & Basson, 2000; Matthew A. Sanders & Basson, 2008; Grande-Allen et al. (manuscript in revision)). Specifically in the context of human enteroids it was observed that changes in a substrates mechanical properties results in differences in growth and viability, in one case even the ability for pathogens to adhere to enteroid (Gjorevski et al., 2016; Hernandez-Gordillo et al., 2020; Ng, Tan, Pek, Tan, & Kurisawa, 2019; Grande-Allen et al. (manuscript in revision)). Substrate stiffness has been reported to affect cell migration and proliferation through the integrin mediated focal adhesion kinase (FAK) pathway, with collagen IV being shown to promote FAK dependent pathways in the caco-2 cell line (Du et al., 2016; M. A. Sanders & Basson, 2000; Matthew A. Sanders & Basson, 2008). Collagen IV substrates were (compared to a poly-L-lysine substrate) also found to influence migration and proliferation of cells through FAK-independent pathways (M. A. Sanders & Basson, 2000; Matthew A. Sanders & Basson, 2008).

Matrigel, an extract derived from a mouse sarcoma line, contains several ECM components including fibronectin, laminin, perlecan, and nidogen-1 (Hughes, Postovit, & Lajoie, 2010; Kleinman & Martin, 2005; Mccarty & Johnson, 2007). Matrigel is commonly used in 3D culture of enteroids due to the ease of polymerizing the matrix at room temperature and depolymerization of the matrix by simply lowering the temperature. However, there have also been reports of successfully using Collagen I as the matrix to grow 3D cultures (Jabaji et al., 2014; Yuli Wang et al., 2017; Yui et al., 2012). Additionally, 2D cultures have more often been grown on a broader number of substrates such as Matrigel, Collagen I, Collagen IV and hydrogels (Hernandez-Gordillo et al., 2020; Jabaji et al., 2013; Lin et al., 2020; Miyoshi & Stappenbeck, 2013; Puzan et al., 2018; Rajan et al., 2020; Sato et al., 2011; Scott et al., 2016; Yuli Wang et al., 2017; Grande-Allen et al. (manuscript in revision)). The components of Matrigel and collagen IV used in the studies explored for this analysis differ in their composition. Thus, the protein attachments to the matrix are likely to be significantly different, and the differential interactions of media components with the matrix are likely to result in altered signaling to the enteroids/colonoids. Furthermore, mechanical stress is likely to be transduced differently by these distinct matrices to the overlying 2D organoid cultures. For example, collagens I and IV vary in their ability to transduce mechanical stress to cells due to differences in how integrins of the cells bind to the substrates (Tang, 2020). In addition to chemical and mechanical forces potentially driving the transcriptional differences between 2D organoids grown on different substrates, there may also be other unknown mechanisms at work, such as the presence of residual growth factors or a yet to be described physical signal. Additionally, it would also be interesting to investigate if the differences that we observed between collagen IV and Matrigel in 2D organoids also exist in 3D organoids.

Here we highlighted one example of a pathway that is influenced by substrate with showing the enrichment of cholesterol homeostats in monolayers on Matrigel in comparison to monolayers on collagen. Absorption of cholesterol by the intestine is a major determinant of cholesterol homeostasis and the target of an important cholesterol-lowering drug, ezetimibe (Malhotra, Gill, Saksena, & Alrefai, 2020; Pradhan, Bhandari, & Sethi, 2020). Furthermore, the cholesterol biosynthesis pathway is essential for homeostasis of the intestinal epithelium (McFarlane et al., 2015; Rong, McDonald, & Engelking, 2017). Thus, a better understanding of cholesterol homeostasis in the intestinal epithelium is important for understanding both cholesterol absorption and epithelial homeostasis.

Together, our results suggest that substrates are not interchangeable and can have a significant influence on the expression of a pathway or gene of interest. It is also important to mention that collagen IV is a basement membrane collagen which is more similar to what the epithelium would be exposed to *in vivo* than Collagen I, which has been used in intestinal organoid 2D and 3D cultures, a highly abundant collagen that is most often found in the bone, skin or tendons. Furthermore, we have provided a resource that can help identify pathways or genes of interest that are enriched in cells grown on these common substrates —we recommend using our current dataset and when designing 2D organoids experiments (Supplementary Information 1, 2 & 3). The format and substrate that the organoids were grown on were the two largest drivers of variance in our dataset. The reason why format is a large driver of variance remains unclear as few investigators have compared organoids grown as 3D, 2D Monolayers and/or Transwells (Guo et al., 2008; Kasendra et al., 2018; Öhlund et al., 2017; T. Wang et al., 2020; Zhang et al., 2017). The differences that we observed between 3D and 2D (monolayers and transwells) formats could be potentially due to the different cell shapes in sheet and spherical conformations, which may alter attachment to the Matrigel substrate. Additionally, the 3D format creates an oxygen gradient with lower oxygen tension in the lumen/apical surface of the organoids, and furthermore requires diffusion of growth factors, nutrients, and cellular metabolites through the matrix; thus, 3D culture creates a niche distinct from 2D culture even when the media and substrate are identical (Okkelman, Neto, Papkovsky, Monaghan, & Dmitriev, 2020).

We also observed variations in the transcriptome of organoids grown in transwells compared to monolayers. While both formats are 2D, significant differences between transwells and monolayers were observed that may be due to transwells allowing exposure of media to both the apical and basolateral surfaces, while in the monolayer cultures the exposure to medium is limited to the apical surface of the cells. For instance, we observed an enrichment in hypoxia signaling in duodenal and jejunal enteroids grown on transwells compared to monolayers, with GSEA. Oxygen levels have been reported to have an effect on the differentiation of pluripotent stem cell derived pancreatic cells with a hypoxic state resulting in a lower amount of differentiated beta cells (Heinis et al., 2010). Changes in metabolism have also been linked with changes in the availably of oxygen. Kondo et al. reported increased oxidative phosphorylation with an increase in oxygen levels (decreased thickness in the culture medium layer) (Kondo et al., 1997). It was reported that under standard cell culture conditions caco-2 monolayers had an oxygen concentration at ∼18% and transwell inserts were reported to have oxygen levels at ∼16% (Fofanova et al., 2019; von Köckritz-Blickwede, Zeitouni, Fandrey, & Naim, 2015). Another study examined the differences between transwells, and air-liquid interface (ALI) cultures based on the diffusion of oxygen into the growth media. They found that ALI cultures, which had a smaller distance between the cell layer and the top of the medium layer, had lower levels of HIF1α because of the increased level of oxygen to the cultures. Furthermore, they reported that ALI cultures had lower HIF1α target genes expressed, and a metabolic shift from producing lactate and performing glycolysis to oxidative phosphorylation (Klasvogt et al., 2017). The availability of oxygen has been shown to cause changes in cells’ differentiation capacity as well as their capacity to shift metabolic pathways. Another reported response to hypoxia is the increase in the production of proinflammatory cytokines at baseline and in response to stimuli (Choi et al., 2017; Fujii et al., 2020; Hsu, Lin, Chiu, Liu, & Huang, 2020; Tannahill et al., 2013). Likewise, we observed an enrichment in interlukin-1 signaling pathways using GSEA of duodenal and jejunal enteroids grown on transwells compared to monolayers. Thus, we believe that the enrichment in the interleukin-1 proinflammatory pathway in transwells may be caused by the hypoxic environment compared to that of monolayers. Furthermore, we have provided a resource that can help identify if a pathway or gene of interest is enriched in a particular format (Supplementary Information 3). To understand the of impact of tissue origin on enteroids in specific culture format we performed unsupervised clustering analyses of the gene expression data. PCA and dendrograms reflected the physical proximity of the segments, where colonoids and ileal enteroids clustered together and were separated from duodenal and jejunal enteroids, which also clustered together (Figures 1, 4A-B, S6). An exception to this was observed in jejunal enteroids transduced with NGN3-expressing lentivirus, which formed a separate cluster even when uninduced, either due to the effect of transduction, leaky expression of *NGN3,* or the Matrigel substrate that the enteroids were grown on (Figure 4A-B). Without taking the Matrigel substrate into account this highlights the importance of when comparing transduced samples to a control, that control should also be a transduced enteroid line to account for these confounding factors such as leaky expression and the influence of transduction on the transcriptome. Ileal enteroids were acquired from the terminal ileum, and the majority of colonic enteroids were derived from ascending colon, therefore it is possible that the physical proximity of the donor tissues may explain the similarity in gene expression as shown by their clustering on PCAs and dendrograms (Foley, O’Flaherty, Barrangou, & Theriot, 2019; Tian et al., 2020; Vuik et al., 2019). A transcriptomic analysis of mouse epithelium and enteroids found a similar segment-specific gene expression pattern but did not include colonoids in their analysis (Middendorp et al., 2014). Additional analyses of human duodenal and ileal enteroids showed that segment-specific markers such as *ASBT* (*SLC10A2*) and *OSTα/β* (*SLC51A/B*) were differentially expressed only in differentiated enteroids (Middendorp et al., 2014). We also observed the differential expression of these genes when comparing our duodenal and ileal enteroids (Supplemental Data 2).

To further understand these gene expression patterns, we used GSEA analysis to identify pathways enriched in distal intestinal organoids compared to proximal organoids. The results for intersegmental comparisons within specific culture formats revealed novel enriched terms as well as those that confirm known functional differences between segments (LaPointe et al., 2008; Mach et al., 2014; Uhlen et al., 2010; Yalong Wang et al., 2020). It is well known that bile acids are released into the duodenum and are reabsorbed by the enterocytes in the ileum and recirculated back to the liver via enterohepatic circulation. The bile acids that escape into the colon are either reabsorbed or interact with the colonic micro-environment and undergo chemical modification by the colonic microbiome (Islam et al., 2011; Molinero, Ruiz, Sánchez, Margolles, & Delgado, 2019; Ridlon, Kang, & Hylemon, 2006). ASBT (SLC10A2) is responsible for the transport of bile acids from the lumen into enterocytes. Subsequently, bile acids are exported out of the basolateral side of enterocytes via OSTα (SLC51A) and OSTβ (SLC51B) (Islam et al., 2011; Molinero et al., 2019; Ridlon et al., 2006). Additionally, Farnesoid X receptor (FXR; NR1H4), a nuclear receptor, is activated by bile acids and promotes the reuptake of bile acids from the intestine to the liver (Matsubara, Li, & Gonzalez, 2013).As expected, this analysis showed an enrichment in the bile acid and salt transport gene set in the ileal enteroids and colonoids relative to the duodenal enteroids (Subramanian et al., 2005). As expected, the segment that organoids are derived from have been shown to maintain their regional transcriptional patterning *in vitro.* This is advantageous because it allows ones to study regional specific responses to specific stimuli over other immortalized or cancer cell lines. For instance, if one is studying the effect of a drug on iron absorption using colonoids or a colon cancer cell line would not accurately recapitulate what would occur in the duodenum where iron is absorbed. Therefore, it is important to take into account the specific region being studied when using a specific intestinal organoid line. We also further investigated the effect of differentiation status on 3D duodenal and colonic organoids. PCA plots, dendrograms and DESeq2 differential gene expression analyses showed the strong effect differentiation media conditions had on 3D enteroids. We found that in dendrograms and PCA plots there was a bifurcation of the samples into the differentiated and undifferentiated groups. We also employed differential gene expression analyses with DESeq2 and found as expected the genes associated with proliferation and stem cells were upregulated in undifferentiated enteroids and genes associated with secretory and absorptive cell types were upregulated in the differentiated organoids (Figure 5 and S7). These results were expected due to the previously mentioned well-studied functional and transcriptional differences between the differentiated and undifferentiated compartments of the intestinal epithelium (Barker, 2014; Middendorp et al., 2014). This also highlights an advantage of human enteroid/colonoid cultures with the ability to separately mimic the crypt or villus compartment of the human intestine. Prior studies using primary cultures, cancer cell lines such as Caco-2 or conditionally immortalized cells were limited in their ability to continuously expand and easily differentiate multiple intestinal epithelial cells as shown in our study (Beaulieu & Ménard, 2012; Perreault & Beaulieu, 1998; Tremblay et al., 2006; Whitehead & Robinson, 2009). Thus, we find that enteroids/colonoids are a facile system for comparing stem/progenitor vs. differentiated cells and can be easily incorporated into the experimental design. For example, it may be preferable to apply luminal stimuli (e.g., food products or infectious agents) to differentiated monolayers or transwell cultures that more closely reflect the apical cell surfaces that first contact these substances. Conversely, to mimic exposure to compounds that will circulate systemically (e.g., chemotherapeutic drugs) and expose the compound to the basal lateral side of the epithelium through the capillaries, it may be important to apply the stimuli to both the differentiated and undifferentiated cultures in either 3D or transwell format. Although our analysis of differentiated versus undifferentiated organoids was focused primarily on 3D cultures, we anticipate that other formats will show similar patterns of gene expression. Therefore, we conclude that the compartment (crypt or villus/upper-differentiated) can be readily incorporated into the experimental design.

An advantage of using enteroids and colonoids over immortalized or cancer cell lines is that organoids more accurately recapitulate normal human intestinal epithelium at homeostasis. Human enteroids and colonoids are also advantageous over inbred mouse strains that lack genetic diversity because with the use of multiple patient derived lines one can more accurately understand the broad response of the human population to a specific stimulus. Therefore, we sought to understand the effect of using enteroid/colonoid lines from various patients on a study. Initially through observing PCA plots and dendrograms we noted that patient samples paired together despite the experimental conditions that the enteroids/colonoids were exposed to (Figure 6 & S8). This phenomenon of patient clustering regardless of experimental manipulation was observed across various segments and formats, indicating that a greater amount of transcriptomic variability exists between different patients than that of experimental conditions such as calcitriol, bacteria and rotavirus infections. It can be observed in the dendrogram for transwells that the batch-to-batch variation in transduced jejunal samples (J2: ULDM1 vs ULDM2) is larger than the variation between technical replicates (J2: ULDM1_1/2/3). And these technical replicates were more similar to each other than 2 different jejunal patient lines from the same segment (Figure 4A&B). To account for this patient-to-patient variability, a paired statistical analysis can be performed; using this approach, we identified 6-7-fold more differentially expressed genes in 3D duodenum and colon organoids than using a standard test of means (Figure 6 &S8). This observation along with the observed patient-to-patient variability that exists in the clinical setting reinforces the need for multiple biological replicates rather than using technical replicates from the same line multiple times. As explored in this manuscript a source of the variation between patients are the variations in their basal gene expression as well as differences in their response to stimuli. Therefore, we recommend using multiple patient lines in human organoid experiments.

Although this study is presented with limitations such as variability in passage numbers, slight variations in the handling and culturing of organoids and unequal datasets, it serves as an excellent tool for understanding the effect of certain variables on the transcriptome of human enteroids/colonoids. We conclude that the enteroid/colonoid system is a powerful surrogate that can serve as an excellent model of the human intestine al epithelium, with the capacity to mimic various segments and compartments. We have found that the culture conditions must be taken into serious consideration when designing experiments, and caution must be used when comparing experiments performed under different experimental conditions. Overall, we highlight the importance of understanding the nuance of *in vitro* model systems and the effect that common variables may have on experimental outcomes with the largest compilation and analysis of human organoids. Finally, our dataset can be used as the basis for rational design of experiments using enteroids/colonoids.

## Methods

### Experimentation

Human intestinal organoids (enteroids and colonoids) data from RNA seq data samples was acquired from multiple labs associated with the Texas Medical Center’s Digestive Disease Core who obtained organoids from the Digestive Disease Consortium Tissue Bank. For all studies, enteroids and colonoids were established from fresh endoscopic biopsies or resected surgical tissue, and maintained as 3D organoid cultures in Matrigel. Some enteroid and colonoid lines were frozen and stored in liquid nitrogen, and subsequently thawed and expanded again as 3D organoid cultures prior to use. Organoids were passaged every 7-14 days and were replated in specific culture conditions below prior to use. Several types of medium were used in the culturing of the enteroids and colonoids used in this study as described previously (Sato et al., 2011; Saxena et al., 2016). CMGF(-) (complete medium without growth factors), consists of advanced DMEM/F-12 medium (gibco) supplemented with 1X GlutaMAX (Invitrogen), 10mM HEPES buffer (gibco), and 100 U/mL penicillin-streptomycin (gibco). CMGF (+) (complete medium with growth factors) is CMGF(-) supplemented with 10% NOGGIN-conditioned medium, 20% R-SPONDIN conditioned medium, 50% Wnt3A-conditioned medium, 1X B-27 supplement (gibco), 1X N-2 supplement (gibco), 10mM nicotinamide (Sigma-Aldrich), 1 mM N-acetylcysteine (Sigma-Aldrich), 10 nM human gastrin I (Sigma-Aldrich), 500 nM A 83-01 (Sigma-Aldrich), 10 µM SB202190 (Sigma-Aldrich) and 50 ng/ml epidermal growth factor (EGF) (R&D Systems). Differentiation medium is CMGF(+) without WNT3A conditioned-medium, nicotinamide and SB202190, and reduced NOGGIN (5%) and R-SPONDIN (10%) conditioned mediums. These mediums were prepared by the Texas Medical Center’s Digestive Disease Core.

In projects 1 and 4, 3D enteroids were generated, maintained and experimentally manipulated in CMGF(+) as previously described (Li et al., 2020). Briefly, undifferentiated and differentiated enteroids/colonoids were grown in CMGF (+) and differentiation medium respectively for 3 days, then treated with equal volumes of calcitriol (100nM) or control (0nM (ethanol)). 24 hours following treatment enteroids/colonoids were collected and RNA isolated with the E.Z.N.A.® Total RNA Kit I (omega BIO-TEK). Paired-end Illumina sequencing libraries were prepared by Novogene (Sacramento, CA, USA). The mRNA sequencing was performed using Illumina platforms for 150 bp paired-end reads. The RNA-seq data (GSE159811) were deposited in the Gene Expression omnibus of the National Center for Biotechnology.

In project 2, enteroids/colonoids from the various intestinal segments were grown and maintained in CMGF(+). After 5-7 days of culture in Matrigel (BD Biosciences) the organoids were dissociated and RNA was isolated with the Qiagen RNeasy kit (Qiagen, Germantown, MD, USA). Paired-end Illumina sequencing libraries were prepared and total RNA-seq was performed with the Hi-seq 2500 (Illumina Inc.).

In project 3, enteroids/colonoids on 96 well plate was generated as previously published (Poole, Rajan, & Maresso, 2018). The undifferentiated enteroids/colonoids monolayers were grown in CMGF(+) for 3 days and in differentiation media from days 3-5. The differentiated monolayers were then infected with bacterial cultures at a multiplicity of infection of 10 for 3 hours. At the end of infection, the cells were lysed in TRIzol (Invitrogen, Waltham, MA, USA) and stored at -80. RNA isolated and prepared for paired-end Illumina sequencing by Novogene (Sacramento, CA, USA).

In project 5, enteroids/colonic were plated on collagen-coated transwell membranes (Corning) differentiated for 5 days following established protocols (Zou et al., 2019). Monolayers on transwells were mock-inoculated with TNC (10 mm Tris-HCl,140 mM NaCl, 10 mM CaCl2, pH 7.4) or inoculated with purified triple-layered human rotavirus (Ito strain) in TNC at a high multiplicity of infection (Saxena et al., PNAS 2017). After 2h of virus adsorption in the presence of 0.2 mg/ml of porcine pancreatin (Sigma-Aldrich) prepared in CMGF(-), HIEs were washed twice with CMGF(-) medium, and then differentiation media containing 0.2 mg/ml of porcine pancreatin was added to the transwell. Two (mock) and four (Ito-infected) transwell membranes per experimental treatment and time point were circumferentially excised and total RNA was extracted immediately using the Qiagen RNeasy Mini Kit. Undifferentiated jejunal 3D organoids from two lines grown in CMGF(+) and similarly harvested. cDNA libraries were prepared using Illumina Epidemiology RiboZero rRNA removal with TruSeq Stranded RNA library prep following the provided protocol (Illumina). Sequencing of the cDNA libraries was performed on a high output v4 flow cell (Illumina) using a paired-end 100 cycle run on a HiSeq 2500 Sequencing System (Illumina).

In project 6, monolayer cultures were prepared as previously described (Ettayebi et al., 2021). Briefly, cell pellets resulting from dispersion of 3D HIEs, were suspended in Intesticult (INT) human organoid growth medium (Stem Cell Technologies) proliferation medium, prepared by mixing equal volumes of components A and B of INT human organoid growth medium, and supplemented with 10 μM ROCK inhibitor Y-27632. After 1 day of cell growth as a monolayer, the proliferation medium was changed with differentiation medium, consisting of an equal volume of component A of INT human organoid growth medium and CMGF(−). The cell monolayers were differentiated for 5 days as previously described. Five-day-differentiated HIE cell monolayers were washed once with CMGF(−)and either mock-infected or inoculated with 5 μl human norovirus diluted in 100 μl CMGF(−) medium supplemented with 500 μM glycochenodeoxycholic acid (GCDCA), for 1 to 2 h at 37°C. The inoculum was removed, and monolayers were washed three times with CMGF(−) to remove unbound virus. Differentiation medium containing 500 μM GCDCA was then added, and the cultures were incubated at 37°C for 24 h. Total RNA was extracted using the Qiagen RNeasy kit and paired-end Illumina sequencing was performed by Novogene (South Plainsfield, NJ, USA).

In project 7, 3D jejunal enteroids were maintained in GMCF(+) as previously described (Saxena et al., 2016). 3D cultures were differentiated for 3-4 days with differentiation medium before inoculation, and were mock-inoculated with TNC or inoculated with purified triple-layered human rotavirus (Ito strain) in TNC at a high multiplicity of infection (Saxena et al., 2017). The HIEs were vigorously pipetted 10 to 20 times with a P200 pipette to disperse and open the HIEs for apical exposure to the virus. After 2h of virus adsorption in the presence of 0.1 to 0.2 mg/ml of porcine pancreatin (Sigma-Aldrich) prepared in CMGF(-) medium, HIEs were washed twice with CMGF(-) medium and centrifuged at 50g to 70g to remove the inoculum and pancreatin that were present during virus adsorption. RNA was extracted using the Qiagen RNeasy Mini kit. Total RNA was prepared by using the Illumina TruSeq Stranded RNA Sample preparation protocol. Paired-end sequencing was performed by using the Illumina HiSeq 2500 machine. The RNA-seq data (GSE90796) were deposited in the Gene Expression omnibus of the National Center for Biotechnology.

In project 8, 3D enteroids initially maintained as 3D cultures in CMGF(+) were disassociated in trypsin/EDTA and seeded onto collagen IV-coated 96 well plates as monolayers. Following culture in CMGF(+) for one day, monolayers were cultured in differentiation medium for 5 days and then mock-inoculated or inoculated with a pandemic GII.4 Syndey strain of human norovirus as a 10% stool filtrate containing a high MOI of virus (1.8X10^6^ genome equivalents/well) or a similar amount of gamma-irradiated stool. After a 1 hr adsorption period, the HIEs were washed twice with CMGF(-) medium to remove unbound viruses. The cultures were harvested at 6, 10 and 24 hours post infection (hpi) for RNA-Seq analyses. Total RNA was extracted from 5-day differentiated, mock-inoculated monolayers at 3 hpi (treated +/- GCDCA for 3 h), and GII.4- or gGII.4-inoculated monolayers at 6, 10 and 24 hpi using the Qiagen RNeasy mini Kit. Libraries were subsequently created and sequencing was performed on an IlluminaHiSeq 2500. The RNA-seq data (GSE150918) were deposited in the Gene Expression omnibus of the National Center for Biotechnology.

In project 9, 3D enteroids were established from human jejunal and duodenal epithelium and cultured as previously described (Saxena et al., 2016). Monolayer cultures of enteroids were cultured atop crosslinked poly(ethylene glycol)-based hydrogels that varied in stiffness as described in Swaminathan et al. (submitted). Briefly, 3D enteroids maintained in CMGF(+) were dissociated into single cells using trypsin, and seeded atop Matrigel-coated hydrogel surfaces at a density of 3x10^5^ cells/cm^2^ for 1 day to form monolayers. After the cells were adhered to the hydrogels, there were grown in differentiation medium for 5 days with medium changes every 48 h. Subsequently, enteroids monolayers harvested in TRIzol, homogenized, and stored at -80°C before being shipped for purification and mRNA-sequencing by Novogene (Sacramento, CA, USA).

In project 10, enteroids on transwell were generated and experimentally manipulated as previously published (Chang-Graham 2019). Briefly, J2 enteroids, which underwent lentivirus transduction in order to integrate into the genome a doxycycline-inducible NGN3 cassette, were seeded onto 24 well Transwells coated with Matrigel and differentiated without doxycycline. After 5 days, differentiation medium from the apical side was removed and enteroids were treated with 100 µL of lyophilized LDM4 that had been resuspended in enteroid differentiation medium (Engevik et al., 2019). After a 3-hour incubation at 37°C, the Transwells were placed into TRIzol and RNA was extracted and purified with a Qiagen RNeasy kit. The RNA-seq data (GSE138350) were deposited in the Gene Expression omnibus of the National Center for Biotechnology.

### RNA-Seq and statistical Analysis

For global assessment of RNA levels from human enteroids samples, Kallsito (version 46.1) was utilized to align RNA-seq libraries to the human reference genome (hg38) and quantify the abundance of those transcripts (Bray, Pimentel, Melsted, & Pachter, 2016). The tximport (version 1.18.0) package was run in R (version 4.0) to create gene-level count matrices for use with DESeq2 (version 1.30) by importing quantification data obtained from Kallisto (Love et al., 2014; Soneson, Love, & Robinson, 2016). DESeq2 was then used to generate transcript levels in each tissue sample. PERMANOVA was performed in R using the adonis function in the vegan package with 999 permutations and pairwise distances were calculated using the Euclidean method. The model used included Line, Status, Segment, Format and Substrate.

Gene set enrichment analysis (GSEA) was used for pathway analysis of gene expression data to find enriched canonical pathways, hallmark pathways, and gene ontology terms as defined by MiSigDB, http://www.broadinstitute.org/gsea/msigdb/index.jsp (Mootha et al., 2003; Subramanian et al., 2005).

Principal component analysis (plotPCA) and volcano plots (EnhancedVolcano) were all generated in R. Dendrograms were generated using hierarchical clustering in the web tool version of iDEP (version 0.91) (Ge, Son, & Yao, 2018). Venn diagrams were generated using web-based tool Venny 2.1 (https://bioinfogp.cnb.csic.es/tools/venny/index.html).

## Supporting information

All Supplemental Tables and Figures

## Acknowledgments

Z Criss was funded by a Howard Hughes Medical Institute Gilliam Fellowship. We thank Dr. William Lagor and Alexandria Doerfler for their insight on the cholesterol biosynthesis pathway. Research reported in this publication was supported by National Institute of Diabetes and Digestive and Kidney Disorders (NIDDK) and National Institute of Allergy and Infectious Diseases (NIAID) of the National Institutes of Health under grant numbers P30DK056338, R01DK118904, U01DK103117, R01DK112365 and U19AI116497. The content is solely the responsibility of the authors and does not necessarily represent the official views of the National Institutes of Health. We are also thankful for supplementary funding provided by Baylor Research Advocates for Student Scientist (BRASS) during the completion of this project. Figure 1 was created with BioRender.com.

## Endnotes

Private sharing link for Figshare data:

Supplementary Information 1: https://figshare.com/s/23bb6b82c72d54592d89

Supplementary Information 2: https://figshare.com/s/13ae9cb8adb499139898

Supplementary Information 3: https://figshare.com/s/101625686647404db374

