## Supplementary material for "Drivers of Transcriptional Variance in Human Intestinal Epithelial Organoids": All Supplemental Tables and Figures

**Supplementary Table 1**

A.)

| ids | Status | Segment | Treatment | Format | Substrate | Project # |
|---|---|---|---|---|---|---|
| C103_D0 | Differentiated | Colon | Control | 3D | Matrigel | 1 |
| C103_D100 | Differentiated | Colon | 1,25D3 | 3D | Matrigel | 1 |
| C104_D0 | Differentiated | Colon | Control | 3D | Matrigel | 1 |
| C104_D100 | Differentiated | Colon | 1,25D3 | 3D | Matrigel | 1 |
| C109_D0 | Differentiated | Colon | Control | 3D | Matrigel | 1 |
| C109_D100 | Differentiated | Colon | 1,25D3 | 3D | Matrigel | 1 |
| C111_D0 | Differentiated | Colon | Control | 3D | Matrigel | 1 |
| C111_D100 | Differentiated | Colon | 1,25D3 | 3D | Matrigel | 1 |
| C112_D0 | Differentiated | Colon | Control | 3D | Matrigel | 1 |
| C112_D100 | Differentiated | Colon | 1,25D3 | 3D | Matrigel | 1 |
| C131_D0 | Differentiated | Colon | Control | 3D | Matrigel | 1 |
| C131_D100 | Differentiated | Colon | 1,25D3 | 3D | Matrigel | 1 |
| C103_H0 | Undifferentiated | Colon | Control | 3D | Matrigel | 1 |
| C103_H100 | Undifferentiated | Colon | 1,25D3 | 3D | Matrigel | 1 |
| C104_H0 | Undifferentiated | Colon | Control | 3D | Matrigel | 1 |
| C104_H100 | Undifferentiated | Colon | 1,25D3 | 3D | Matrigel | 1 |
| C109_H0 | Undifferentiated | Colon | Control | 3D | Matrigel | 1 |
| C109_H100 | Undifferentiated | Colon | 1,25D3 | 3D | Matrigel | 1 |
| C111_H0 | Undifferentiated | Colon | Control | 3D | Matrigel | 1 |
| C111_H100 | Undifferentiated | Colon | 1,25D3 | 3D | Matrigel | 1 |
| C112_H0 | Undifferentiated | Colon | Control | 3D | Matrigel | 1 |
| C112_H100 | Undifferentiated | Colon | 1,25D3 | 3D | Matrigel | 1 |
| C131_H0 | Undifferentiated | Colon | Control | 3D | Matrigel | 1 |
| C131_H100 | Undifferentiated | Colon | 1,25D3 | 3D | Matrigel | 1 |
| D102_D0 | Differentiated | Duodenum | Control | 3D | Matrigel | 1 |
| D102_D100 | Differentiated | Duodenum | 1,25D3 | 3D | Matrigel | 1 |
| D103_D0 | Differentiated | Duodenum | Control | 3D | Matrigel | 1 |
| D103_D100 | Differentiated | Duodenum | 1,25D3 | 3D | Matrigel | 1 |
| D104_D0 | Differentiated | Duodenum | Control | 3D | Matrigel | 1 |
| D104_D100 | Differentiated | Duodenum | 1,25D3 | 3D | Matrigel | 1 |
| D109_D0 | Differentiated | Duodenum | Control | 3D | Matrigel | 1 |
| D109_D100 | Differentiated | Duodenum | 1,25D3 | 3D | Matrigel | 1 |
| D144_D0 | Differentiated | Duodenum | Control | 3D | Matrigel | 1 |
| D144_D100 | Differentiated | Duodenum | 1,25D3 | 3D | Matrigel | 1 |
| D2_D0 | Differentiated | Duodenum | Control | 3D | Matrigel | 1 |
| D2_D100 | Differentiated | Duodenum | 1,25D3 | 3D | Matrigel | 1 |
| D102_H0 | Undifferentiated | Duodenum | Control | 3D | Matrigel | 1 |
| D102_H100 | Undifferentiated | Duodenum | 1,25D3 | 3D | Matrigel | 1 |
| D103_H0 | Undifferentiated | Duodenum | Control | 3D | Matrigel | 1 |
| D103_H100 | Undifferentiated | Duodenum | 1,25D3 | 3D | Matrigel | 1 |
| D104_H0 | Undifferentiated | Duodenum | Control | 3D | Matrigel | 1 |
| D104_H100 | Undifferentiated | Duodenum | 1,25D3 | 3D | Matrigel | 1 |
| D109_H0 | Undifferentiated | Duodenum | Control | 3D | Matrigel | 1 |
| D109_H100 | Undifferentiated | Duodenum | 1,25D3 | 3D | Matrigel | 1 |
| D144_H0 | Undifferentiated | Duodenum | Control | 3D | Matrigel | 1 |
| D144_H100 | Undifferentiated | Duodenum | 1,25D3 | 3D | Matrigel | 1 |
| D2_H0 | Undifferentiated | Duodenum | Control | 3D | Matrigel | 1 |
| D2_H100 | Undifferentiated | Duodenum | 1,25D3 | 3D | Matrigel | 1 |
| C143 | Undifferentiated | Colon | Control | 3D | Matrigel | 2 |
| C144 | Undifferentiated | Colon | Control | 3D | Matrigel | 2 |
| C146 | Undifferentiated | Colon | Control | 3D | Matrigel | 2 |
| C147 | Undifferentiated | Colon | Control | 3D | Matrigel | 2 |
| C152 | Undifferentiated | Colon | Control | 3D | Matrigel | 2 |
| C153 | Undifferentiated | Colon | Control | 3D | Matrigel | 2 |
| C173_CD | Undifferentiated | Colon | Control | 3D | Matrigel | 2 |
| C177_CD | Undifferentiated | Colon | Control | 3D | Matrigel | 2 |
| C183_CD | Undifferentiated | Colon | Control | 3D | Matrigel | 2 |
| C189_CD | Undifferentiated | Colon | Control | 3D | Matrigel | 2 |
| C203_CD | Undifferentiated | Colon | Control | 3D | Matrigel | 2 |
| C209_CD | Undifferentiated | Colon | Control | 3D | Matrigel | 2 |
| C3_aafA | Differentiated | Colon | aafA | Monolayer | Matrigel | 3 |
| C3_eaec | Differentiated | Colon | eaec | Monolayer | Matrigel | 3 |
| C3_ehec | Differentiated | Colon | ehec | Monolayer | Matrigel | 3 |
| C3_expec | Differentiated | Colon | expec | Monolayer | Matrigel | 3 |
| C3_HS | Differentiated | Colon | HS | Monolayer | Matrigel | 3 |
| C3_Mock | Differentiated | Colon | Mock | Monolayer | Matrigel | 3 |
| C4_aafA | Differentiated | Colon | aafA | Monolayer | Matrigel | 3 |
| c4_eaec | Differentiated | Colon | eaec | Monolayer | Matrigel | 3 |
| C4_ehec | Differentiated | Colon | ehec | Monolayer | Matrigel | 3 |
| C4_expec | Differentiated | Colon | expec | Monolayer | Matrigel | 3 |
| C4_HS | Differentiated | Colon | HS | Monolayer | Matrigel | 3 |
| C4_Mock | Differentiated | Colon | Mock | Monolayer | Matrigel | 3 |
| C9_aafA | Differentiated | Colon | aafA | Monolayer | Matrigel | 3 |
| C9_eaec | Differentiated | Colon | eaec | Monolayer | Matrigel | 3 |
| C9_ehec | Differentiated | Colon | ehec | Monolayer | Matrigel | 3 |
| C9_expec | Differentiated | Colon | expec | Monolayer | Matrigel | 3 |
| C9_HS | Differentiated | Colon | HS | Monolayer | Matrigel | 3 |
| C9_Mock | Differentiated | Colon | Mock | Monolayer | Matrigel | 3 |
| D103_aafA | Differentiated | Duodenum | aafA | Monolayer | Matrigel | 3 |
| D103_HS | Differentiated | Duodenum | HS | Monolayer | Matrigel | 3 |
| D103_Mock | Differentiated | Duodenum | Mock | Monolayer | Matrigel | 3 |
| D3_eaec | Differentiated | Duodenum | eaec | Monolayer | Matrigel | 3 |
| D3_ehec | Differentiated | Duodenum | ehec | Monolayer | Matrigel | 3 |
| D3_expec | Differentiated | Duodenum | expec | Monolayer | Matrigel | 3 |

**Supplementary Table 1**

| | | | | | | |
|---|---|---|---|---|---|---|
| D4_aafA | Differentiated | Duodenum | aafA | Monolayer | Matrigel | 3 |
| D4_eaec | Differentiated | Duodenum | eaec | Monolayer | Matrigel | 3 |
| D4_ehec | Differentiated | Duodenum | ehec | Monolayer | Matrigel | 3 |
| D4_expec | Differentiated | Duodenum | expec | Monolayer | Matrigel | 3 |
| D4_HS | Differentiated | Duodenum | HS | Monolayer | Matrigel | 3 |
| D4_Mock | Differentiated | Duodenum | Mock | Monolayer | Matrigel | 3 |
| D9_aafA | Differentiated | Duodenum | aafA | Monolayer | Matrigel | 3 |
| D9_eaec | Differentiated | Duodenum | eaec | Monolayer | Matrigel | 3 |
| D9_ehec | Differentiated | Duodenum | ehec | Monolayer | Matrigel | 3 |
| D9_expec | Differentiated | Duodenum | expec | Monolayer | Matrigel | 3 |
| D9_HS | Differentiated | Duodenum | HS | Monolayer | Matrigel | 3 |
| D9_Mock | Differentiated | Duodenum | Mock | Monolayer | Matrigel | 3 |
| I103_aafA | Differentiated | Ileum | aafA | Monolayer | Matrigel | 3 |
| I103_ehec | Differentiated | Ileum | ehec | Monolayer | Matrigel | 3 |
| I103_HS | Differentiated | Ileum | HS | Monolayer | Matrigel | 3 |
| I103_Mock | Differentiated | Ileum | Mock | Monolayer | Matrigel | 3 |
| I3_eaec | Differentiated | Ileum | eaec | Monolayer | Matrigel | 3 |
| I3_expec | Differentiated | Ileum | expec | Monolayer | Matrigel | 3 |
| I4_aafA | Differentiated | Ileum | aafA | Monolayer | Matrigel | 3 |
| I4_eaec | Differentiated | Ileum | eaec | Monolayer | Matrigel | 3 |
| I4_ehec | Differentiated | Ileum | ehec | Monolayer | Matrigel | 3 |
| I4_expec | Differentiated | Ileum | expec | Monolayer | Matrigel | 3 |
| I4_HS | Differentiated | Ileum | HS | Monolayer | Matrigel | 3 |
| I4_Mock | Differentiated | Ileum | Mock | Monolayer | Matrigel | 3 |
| I9_aafA | Differentiated | Ileum | aafA | Monolayer | Matrigel | 3 |
| I9_eaec | Differentiated | Ileum | eaec | Monolayer | Matrigel | 3 |
| I9_ehec | Differentiated | Ileum | ehec | Monolayer | Matrigel | 3 |
| I9_expec | Differentiated | Ileum | expec | Monolayer | Matrigel | 3 |
| I9_HS | Differentiated | Ileum | HS | Monolayer | Matrigel | 3 |
| J11_aafA | Differentiated | Jejunum | aafA | Monolayer | Matrigel | 3 |
| J11_eaec | Differentiated | Jejunum | eaec | Monolayer | Matrigel | 3 |
| J11_ehec | Differentiated | Jejunum | ehec | Monolayer | Matrigel | 3 |
| J11_expec | Differentiated | Jejunum | expec | Monolayer | Matrigel | 3 |
| J11_HS | Differentiated | Jejunum | HS | Monolayer | Matrigel | 3 |
| J11_Mock | Differentiated | Jejunum | Mock | Monolayer | Matrigel | 3 |
| J2_aafA | Differentiated | Jejunum | aafA | Monolayer | Matrigel | 3 |
| J2_eaec | Differentiated | Jejunum | eaec | Monolayer | Matrigel | 3 |
| J2_ehec | Differentiated | Jejunum | ehec | Monolayer | Matrigel | 3 |
| J2_expec | Differentiated | Jejunum | expec | Monolayer | Matrigel | 3 |
| J2_HS | Differentiated | Jejunum | HS | Monolayer | Matrigel | 3 |
| J2_Mock | Differentiated | Jejunum | Mock | Monolayer | Matrigel | 3 |
| J3_aafA | Differentiated | Jejunum | aafA | Monolayer | Matrigel | 3 |
| J3_eaec | Differentiated | Jejunum | eaec | Monolayer | Matrigel | 3 |
| J3_ehec | Differentiated | Jejunum | ehec | Monolayer | Matrigel | 3 |
| J3_expec | Differentiated | Jejunum | expec | Monolayer | Matrigel | 3 |
| J3_HS | Differentiated | Jejunum | HS | Monolayer | Matrigel | 3 |
| J3_Mock | Differentiated | Jejunum | Mock | Monolayer | Matrigel | 3 |
| C135_D0 | Differentiated | Colon | Control | 3D | Matrigel | 4 |
| C136_D0 | Differentiated | Colon | Control | 3D | Matrigel | 4 |
| C136_H0 | Undifferentiated | Colon | Control | 3D | Matrigel | 4 |
| C103_bac_6h | Differentiated | Colon | bac | Transwell | Collagen | 5 |
| C103_Ito_24h | Differentiated | Colon | Ito | Transwell | Collagen | 5 |
| C103_Ito_6h | Differentiated | Colon | Ito | Transwell | Collagen | 5 |
| C103_mock_24h | Differentiated | Colon | Mock | Transwell | Collagen | 5 |
| C103_mock_6h | Differentiated | Colon | Mock | Transwell | Collagen | 5 |
| C103_mock_bac_6h | Differentiated | Colon | Mock-bac | Transwell | Collagen | 5 |
| C109_bac_6h | Differentiated | Colon | bac | Transwell | Collagen | 5 |
| C109_Ito_24h | Differentiated | Colon | Ito | Transwell | Collagen | 5 |
| C109_Ito_6h | Differentiated | Colon | Ito | Transwell | Collagen | 5 |
| C109_mock_24h | Differentiated | Colon | Mock | Transwell | Collagen | 5 |
| C109_mock_6h | Differentiated | Colon | Mock | Transwell | Collagen | 5 |
| C109_mock_bac_6h | Differentiated | Colon | Mock-bac | Transwell | Collagen | 5 |
| D103_bac_6h | Differentiated | Duodenum | bac | Transwell | Collagen | 5 |
| D103_Ito_24h | Differentiated | Duodenum | Ito | Transwell | Collagen | 5 |
| D103_mock_24h | Differentiated | Duodenum | Mock | Transwell | Collagen | 5 |
| D103_mock_6h | Differentiated | Duodenum | Mock | Transwell | Collagen | 5 |
| D103_mock_bac_6h | Differentiated | Duodenum | Mock-bac | Transwell | Collagen | 5 |
| D109_bac_6h | Differentiated | Duodenum | bac | Transwell | Collagen | 5 |
| D109_Ito_24h | Differentiated | Duodenum | Ito | Transwell | Collagen | 5 |
| D109_Ito_6h | Differentiated | Duodenum | Ito | Transwell | Collagen | 5 |
| D109_mock_24h | Differentiated | Duodenum | Mock | Transwell | Collagen | 5 |
| D109_mock_6h | Differentiated | Duodenum | Mock | Transwell | Collagen | 5 |
| D109_mock_bac_6h | Differentiated | Duodenum | Mock-bac | Transwell | Collagen | 5 |
| I103_bac_6h | Differentiated | Ileum | bac | Transwell | Collagen | 5 |
| I103_Ito_24h | Differentiated | Ileum | Ito | Transwell | Collagen | 5 |
| I103_Ito_6h | Differentiated | Ileum | Ito | Transwell | Collagen | 5 |
| I103_mock_24h | Differentiated | Ileum | Mock | Transwell | Collagen | 5 |
| I103_mock_6h | Differentiated | Ileum | Mock | Transwell | Collagen | 5 |
| I103_mock_bac_6h | Differentiated | Ileum | Mock-bac | Transwell | Collagen | 5 |
| I109_bac_6h | Differentiated | Ileum | bac | Transwell | Collagen | 5 |
| I109_Ito_24h | Differentiated | Ileum | Ito | Transwell | Collagen | 5 |
| I109_Ito_6h | Differentiated | Ileum | Ito | Transwell | Collagen | 5 |
| I109_mock_24h | Differentiated | Ileum | Mock | Transwell | Collagen | 5 |
| I109_mock_6h | Differentiated | Ileum | Mock | Transwell | Collagen | 5 |
| I109_mock_bac_6h | Differentiated | Ileum | Mock-bac | Transwell | Collagen | 5 |

**Supplementary Table 1**

| | | | | | | |
|---|---|---|---|---|---|---|
| **J11_bac_6h** | Differentiated | Jejunum | bac | Transwell | Collagen | 5 |
| **J11_Ito_6h** | Differentiated | Jejunum | Ito | Transwell | Collagen | 5 |
| **J11_mock_24h** | Differentiated | Jejunum | Mock | Transwell | Collagen | 5 |
| **J11_mock_6h** | Differentiated | Jejunum | Mock | Transwell | Collagen | 5 |
| **J11_mock_bac_6h** | Differentiated | Jejunum | Mock-bac | Transwell | Collagen | 5 |
| **J2_bac_6h** | Differentiated | Jejunum | bac | Transwell | Collagen | 5 |
| **J2_Ito_24h** | Differentiated | Jejunum | Ito | Transwell | Collagen | 5 |
| **J2_Ito_6h** | Differentiated | Jejunum | Ito | Transwell | Collagen | 5 |
| **J2_mock_24h** | Differentiated | Jejunum | Mock | Transwell | Collagen | 5 |
| **J2_mock_6h** | Differentiated | Jejunum | Mock | Transwell | Collagen | 5 |
| **J2_mock_bac_6h** | Differentiated | Jejunum | Mock-bac | Transwell | Collagen | 5 |
| **D104_MO** | Differentiated | Duodenum | Mock | Monolayer | Collagen | 6 |
| **D104_VO** | Differentiated | Duodenum | Virus | Monolayer | Collagen | 6 |
| **D109_MO** | Differentiated | Duodenum | Mock | Monolayer | Collagen | 6 |
| **D109_VO** | Differentiated | Duodenum | Virus | Monolayer | Collagen | 6 |
| **D4_MO** | Differentiated | Duodenum | Mock | Monolayer | Collagen | 6 |
| **D4_VO** | Differentiated | Duodenum | Virus | Monolayer | Collagen | 6 |
| **D8_MO** | Differentiated | Duodenum | Mock | Monolayer | Collagen | 6 |
| **D8_VO** | Differentiated | Duodenum | Virus | Monolayer | Collagen | 6 |
| **j2_Ito** | Differentiated | Jejunum | Ito | 3D | Matrigel | 7 |
| **j2_mock_** | Differentiated | Jejunum | Mock | 3D | Matrigel | 7 |
| **j7_Ito** | Differentiated | Jejunum | Ito | 3D | Matrigel | 7 |
| **j7_mock** | Differentiated | Jejunum | Mock | 3D | Matrigel | 7 |
| **J11** | Undifferentiated | Jejunum | Control | 3D | Matrigel | 7 |
| **J2** | Undifferentiated | Jejunum | Control | 3D | Matrigel | 7 |
| **J11_GCDCA_3h** | Differentiated | Jejunum | GCDCA | Monolayer | Collagen | 8 |
| **J11_gGII.4_10h** | Differentiated | Jejunum | gGII.4 | Monolayer | Collagen | 8 |
| **J11_gGII.4_24h** | Differentiated | Jejunum | gGII.4 | Monolayer | Collagen | 8 |
| **J11_gGII.4_6h** | Differentiated | Jejunum | gGII.4 | Monolayer | Collagen | 8 |
| **J11_GII.4_10h** | Differentiated | Jejunum | GII.4 | Monolayer | Collagen | 8 |
| **J11_GII.4_24h** | Differentiated | Jejunum | GII.4 | Monolayer | Collagen | 8 |
| **J11_GII.4_6h** | Differentiated | Jejunum | GII.4 | Monolayer | Collagen | 8 |
| **J11_media_3h** | Differentiated | Jejunum | Media | Monolayer | Collagen | 8 |
| **J2_GCDCA_3h** | Differentiated | Jejunum | GCDCA | Monolayer | Collagen | 8 |
| **J2_gGII.4_10h** | Differentiated | Jejunum | gGII.4 | Monolayer | Collagen | 8 |
| **J2_gGII.4_24h** | Differentiated | Jejunum | gGII.4 | Monolayer | Collagen | 8 |
| **J2_gGII.4_6h** | Differentiated | Jejunum | gGII.4 | Monolayer | Collagen | 8 |
| **J2_GII.4_10h** | Differentiated | Jejunum | GII.4 | Monolayer | Collagen | 8 |
| **J2_GII.4_24h** | Differentiated | Jejunum | GII.4 | Monolayer | Collagen | 8 |
| **J2_GII.4_6h** | Differentiated | Jejunum | GII.4 | Monolayer | Collagen | 8 |
| **J2_media_3h** | Differentiated | Jejunum | Media | Monolayer | Collagen | 8 |
| **SOJ11_2** | Differentiated | Jejunum | Soft | Monolayer | Hydrogel | 9 |
| **SOJ11_3** | Differentiated | Jejunum | Soft | Monolayer | Hydrogel | 9 |
| **J2soft5** | Differentiated | Jejunum | Soft | Monolayer | Hydrogel | 9 |
| **J2sof6** | Differentiated | Jejunum | Soft | Monolayer | Hydrogel | 9 |
| **J2sof7** | Differentiated | Jejunum | Soft | Monolayer | Hydrogel | 9 |
| **SOJ3_1** | Differentiated | Jejunum | Soft | Monolayer | Hydrogel | 9 |
| **SOJ3_2** | Differentiated | Jejunum | Soft | Monolayer | Hydrogel | 9 |
| **SOJ3_3** | Differentiated | Jejunum | Soft | Monolayer | Hydrogel | 9 |
| **MJ11_1** | Differentiated | Jejunum | Medium | Monolayer | Hydrogel | 9 |
| **MJ11_2** | Differentiated | Jejunum | Medium | Monolayer | Hydrogel | 9 |
| **MJ11_3** | Differentiated | Jejunum | Medium | Monolayer | Hydrogel | 9 |
| **J2md5** | Differentiated | Jejunum | Medium | Monolayer | Hydrogel | 9 |
| **J2med6** | Differentiated | Jejunum | Medium | Monolayer | Hydrogel | 9 |
| **J2med7** | Differentiated | Jejunum | Medium | Monolayer | Hydrogel | 9 |
| **MJ3_1** | Differentiated | Jejunum | Medium | Monolayer | Hydrogel | 9 |
| **MJ3_2** | Differentiated | Jejunum | Medium | Monolayer | Hydrogel | 9 |
| **MJ3_3** | Differentiated | Jejunum | Medium | Monolayer | Hydrogel | 9 |
| **STJ11_1** | Differentiated | Jejunum | Stiff | Monolayer | Hydrogel | 9 |
| **STJ11_2** | Differentiated | Jejunum | Stiff | Monolayer | Hydrogel | 9 |
| **J11stiff5** | Differentiated | Jejunum | Stiff | Monolayer | Hydrogel | 9 |
| **J2st5** | Differentiated | Jejunum | Stiff | Monolayer | Hydrogel | 9 |
| **J2st6** | Differentiated | Jejunum | Stiff | Monolayer | Hydrogel | 9 |
| **J2st7** | Differentiated | Jejunum | Stiff | Monolayer | Hydrogel | 9 |
| **STJ3_1** | Differentiated | Jejunum | Stiff | Monolayer | Hydrogel | 9 |
| **STJ3_2** | Differentiated | Jejunum | Stiff | Monolayer | Hydrogel | 9 |
| **STJ3_3** | Differentiated | Jejunum | Stiff | Monolayer | Hydrogel | 9 |
| **A96WJ111** | Differentiated | Jejunum | Plastic | Monolayer | Matrigel | 9 |
| **A96WJ112** | Differentiated | Jejunum | Plastic | Monolayer | Matrigel | 9 |
| **A96WJ113** | Differentiated | Jejunum | Plastic | Monolayer | Matrigel | 9 |
| **J2961** | Differentiated | Jejunum | Plastic | Monolayer | Matrigel | 9 |
| **J2962** | Differentiated | Jejunum | Plastic | Monolayer | Matrigel | 9 |
| **J2967** | Differentiated | Jejunum | Plastic | Monolayer | Matrigel | 9 |
| **A96WJ3_1** | Differentiated | Jejunum | Plastic | Monolayer | Matrigel | 9 |
| **A96WJ3_2** | Differentiated | Jejunum | Plastic | Monolayer | Matrigel | 9 |
| **J3967** | Differentiated | Jejunum | Plastic | Monolayer | Matrigel | 9 |
| **ULDM1_1** | Differentiated | Jejunum | LDM4_Uninduced_NGN3 | Transwell | Matrigel | 10 |
| **ULDM1_2** | Differentiated | Jejunum | LDM4_Uninduced_NGN3 | Transwell | Matrigel | 10 |
| **ULDM1_3** | Differentiated | Jejunum | LDM4_Uninduced_NGN3 | Transwell | Matrigel | 10 |
| **ULDM2_1** | Differentiated | Jejunum | LDM4_Uninduced_NGN3 | Transwell | Matrigel | 10 |
| **ULDM2_2** | Differentiated | Jejunum | LDM4_Uninduced_NGN3 | Transwell | Matrigel | 10 |
| **ULDM2_3** | Differentiated | Jejunum | LDM4_Uninduced_NGN3 | Transwell | Matrigel | 10 |

**Supplementary Table 1.) Demographics of data used in analysis - Table of Samples** A.) Shown is the metadata table with the Status, Segment, Treatment, Format, Substrate and Project for every sample (ids) used in this analysis (n=251).

Supplementary Table 2

A.)

| DESeq2 Differential Gene Expression Results (FDR ≤ 0.01 and a fold change ≥2 or ≤0.5) | | |
|---|---|---|
| | **Comparison** | **DE Genes** |
| **Differentiated Duodenum** | 3D-MATRIGEL_vs_MONOLAYER-MATRIGEL | 5139 |
| | MONOLAYER-COLLAGEN_vs_MONOLAYER-MATRIGEL | 6004 |
| | MONOLAYER-COLLAGEN_vs_TRANSWELL-COLLAGEN | 4695 |
| **Differentiated Jejunum** | 3D-MATRIGEL_vs_MONOLAYER-MATRIGEL | 5195 |
| | 3D-MATRIGEL_vs_TRANSWELL-COLLAGEN | 1305 |
| | MONOLAYER-COLLAGEN_vs_MONOLAYER-HYDROGEL | 7113 |
| | MONOLAYER-COLLAGEN_vs_TRANSWELL-COLLAGEN | 3740 |
| | MONOLAYER-MATRIGEL_vs_MONOLAYER-COLLAGEN | 5148 |
| | MONOLAYER-MATRIGEL_vs_MONOLAYER-HYDROGEL | 4341 |
| **Differentiated Ileum** | MONOLAYER_MATRIGEL_vs_TRANSWELL_COLLAGEN | 5702 |
| **Differentiated Colon** | MONOLAYER-MATRIGEL_vs_3D-MATRIGEL | 6049 |
| | MONOLAYER-MATRIGEL_vs_TRANSWELL-COLLAGEN | 5253 |
| | TRANSWELL-COLLAGEN_vs_3D-MATRIGEL | 7271 |
| **Transwells** | DUODENUM_VS_JEJUNUM | 420 |
| | DUODENUM_VS_ILEUM | 3115 |
| | DUODENUM_VS_COLON | 3720 |
| | JEJUNUM_VS_ILEUM | 2857 |
| | JEJUNUM_VS_COLON | 3801 |
| | ILEUM_VS_COLON | 2592 |
| **Monolayers** | DUODENUM_VS_JEJUNUM | 1012 |
| | DUODENUM_VS_ILEUM | 2184 |
| | DUODENUM_VS_COLON | 2750 |
| | JEJUNUM_VS_ILEUM | 3206 |
| | JEJUNUM_VS_COLON | 4115 |
| | ILEUM_VS_COLON | 1266 |
| **3D** | DUODENUM_vs_COLON_DIFFERENTIATED | 2296 |
| | JEJUNUM_vs_COLON_DIFFERENTIATED | 5977 |
| | DUODENUM_vs_JEJUNUM_DIFFERENTIATED | 5332 |
| | DUODENUM_vs_COLON_UNDIFFERENTIATED | 2224 |
| | JEJUNUM_vs_COLON_UNDIFFERENTIATED | 3705 |
| | DUODENUM_vs_JEJUNUM_UNDIFFERENTIATED | 3873 |
| | DUODENUM: DIFFERENTIATED_vs_UNDIFFERENTIATED | 2902 |
| | COLON: DIFFERENTIATED_vs_UNDIFFERENTIATED | 2756 |

**Supplementary Table 2. Differentially expressed genes via DESeq2.** Summary table of the number of differentially expressed genes for 33 comparisons performed in DESeq2 (FDR ≤ 0.01 & Fold Change ≥2 and ≤ 0.5). DE Gene: Differentially Expressed Genes.

**Supplementary Figure 1**

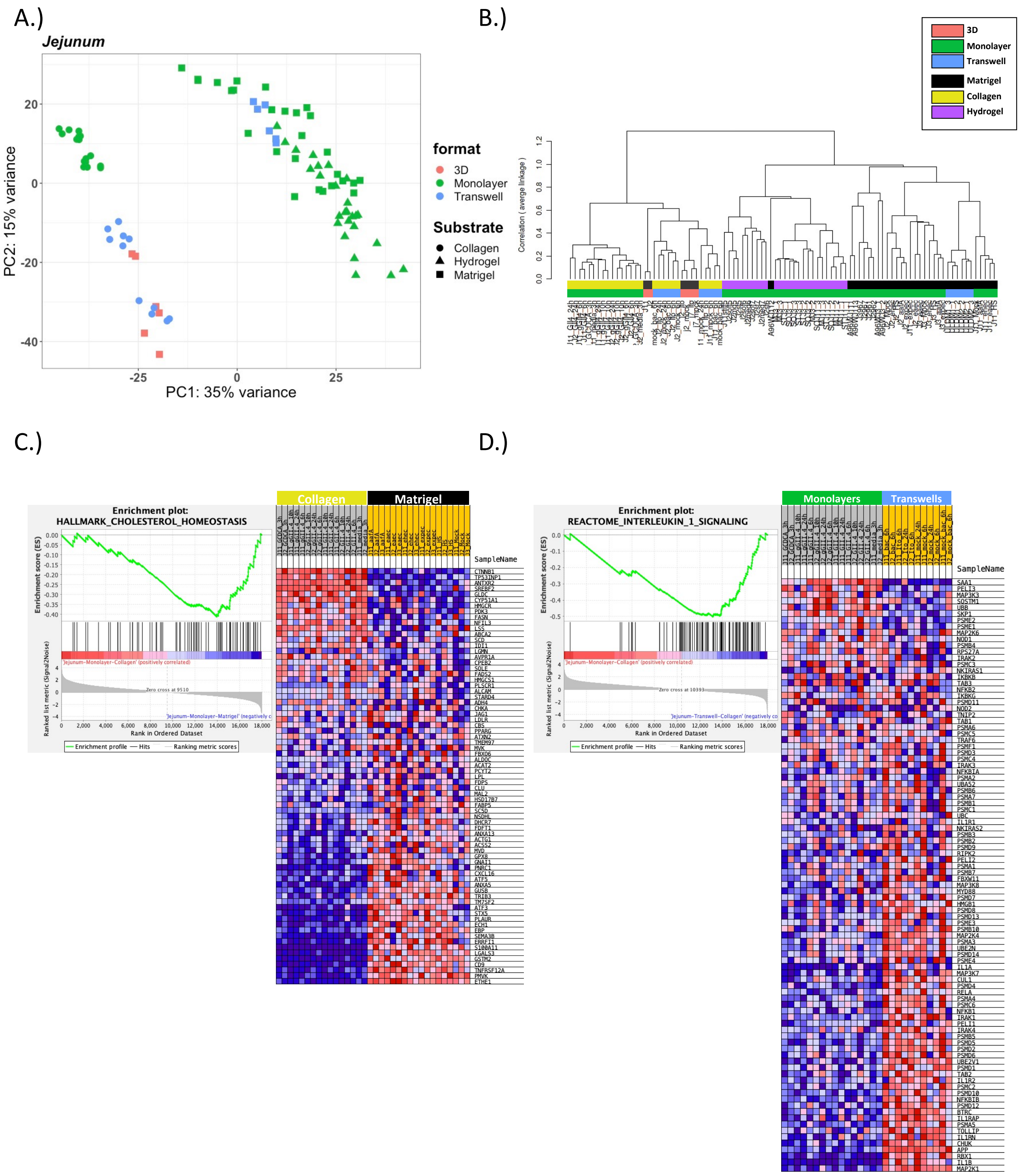

**Supplementary Figure 1. Cholesterol biosynthesis and Interleukin-1 signaling pathway genes are upregulated in jejunal monolayers grown on Matrigel and jejunal transwells grown on collagen, respectively.** A.) PCA of the RNA-sequencing datasets for jejunal enteroids: 3D-Matrigel (n=6), Monolayer-Collagen (n=16), Monolayer-Hydrogel (n=26), Monolayer-Matrigel (n=27), Transwell-Collagen (n=11), and Transwell-Matrigel(n=6); B.) A dendrogram with agglomerative hierarchal clustering of the jejunal gene set from RNA-sequencing. Branch length indicates degree of difference between samples; C.) GSEA showing an enrichment of the hallmark cholesterol homeostasis get set signature in jejunal monolayers grown on Matrigel when compared to jejunal monolayers grown on collagen; D.) GSEA showing an enrichment of the reactome interleukin-1 signaling get set signature in jejunal transwells grown on collagen when compared to jejunal monolayers grown on collagen;

**Supplementary Figure 2**

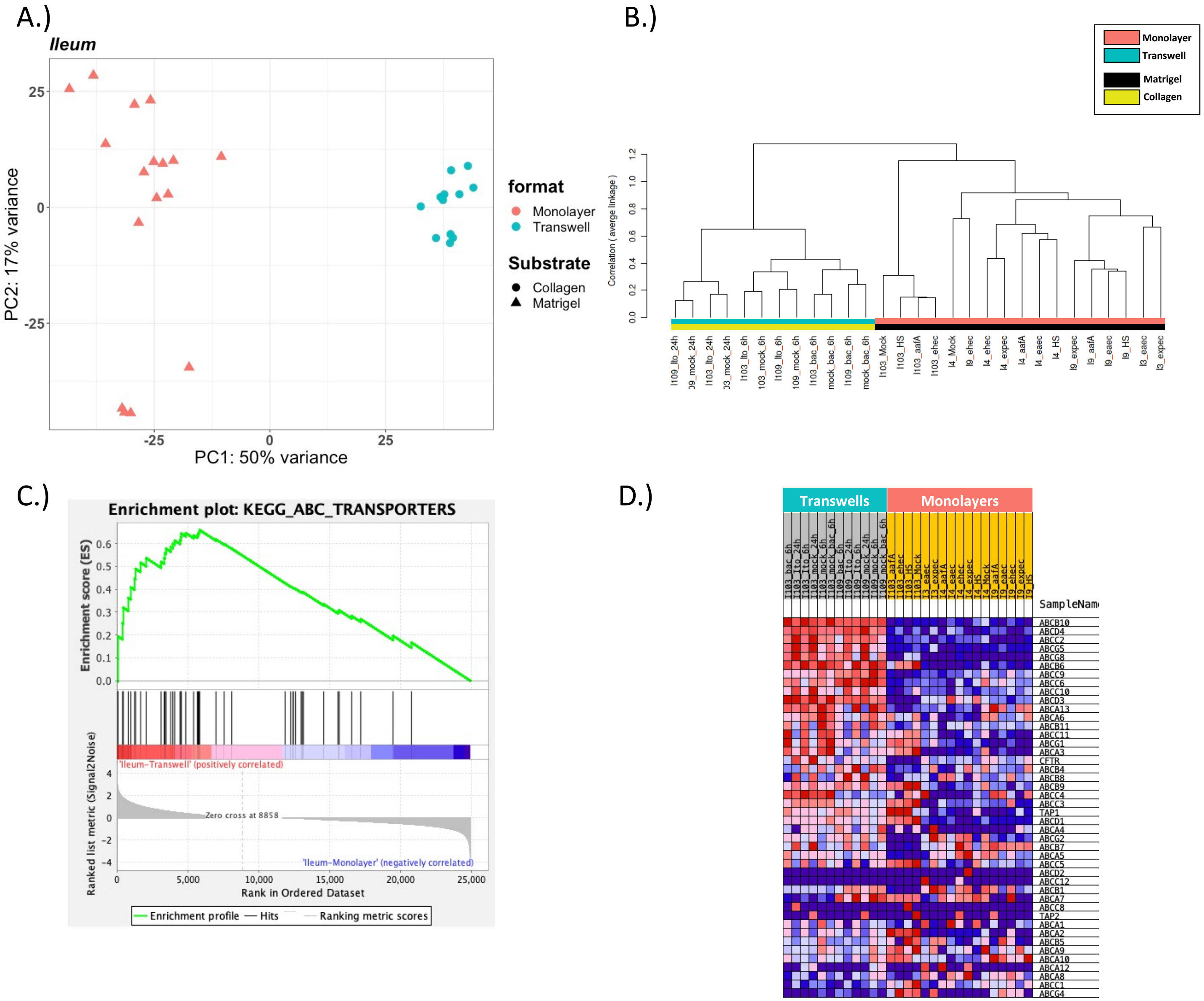


**Supplementary Figure 2. ABC transporter genes are upregulated in Ileal transwells grown on collagen in comparison to ileal monolayers grown on Matrigel.** A.) PCA of the RNA-sequencing datasets for ileal enteroids: Monolayer-Matrigel (n=17), and Transwell-Collagen (n=12); B.) A dendrogram with agglomerative hierarchal clustering of the Ileal gene set from RNA-sequencing. Branch length indicates degree of difference between samples; C.) GSEA showing an enrichment of the KEGG ABC transporters gene set signature in ileal transwells grown on collagen IV when compared to ileal monolayers grown on Matrigel; D.) Heatmap of the KEGG ABC transporters gene set showing an enrichment in ileal transwells grown on collagen IV (gray) compared to ileal monolayers grown on Matrigel (yellow);

**Supplementary Figure 3**

A.)

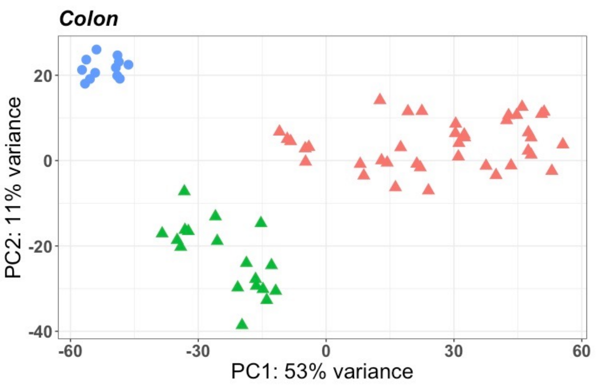


B.)

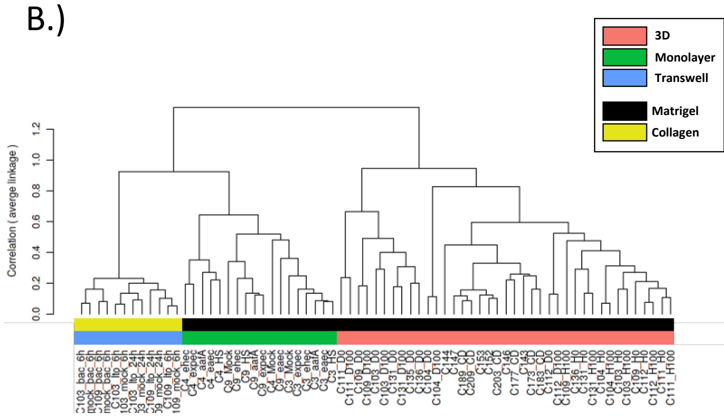


C.)

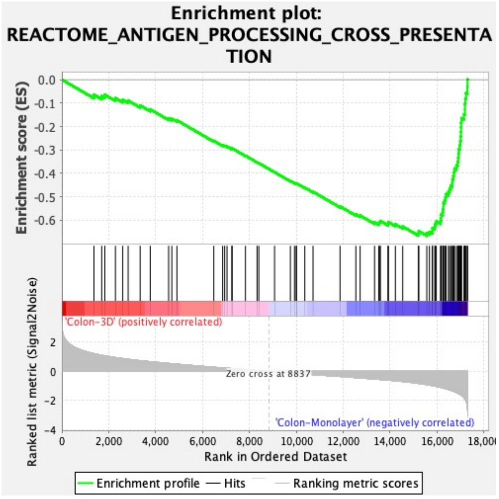


D.)

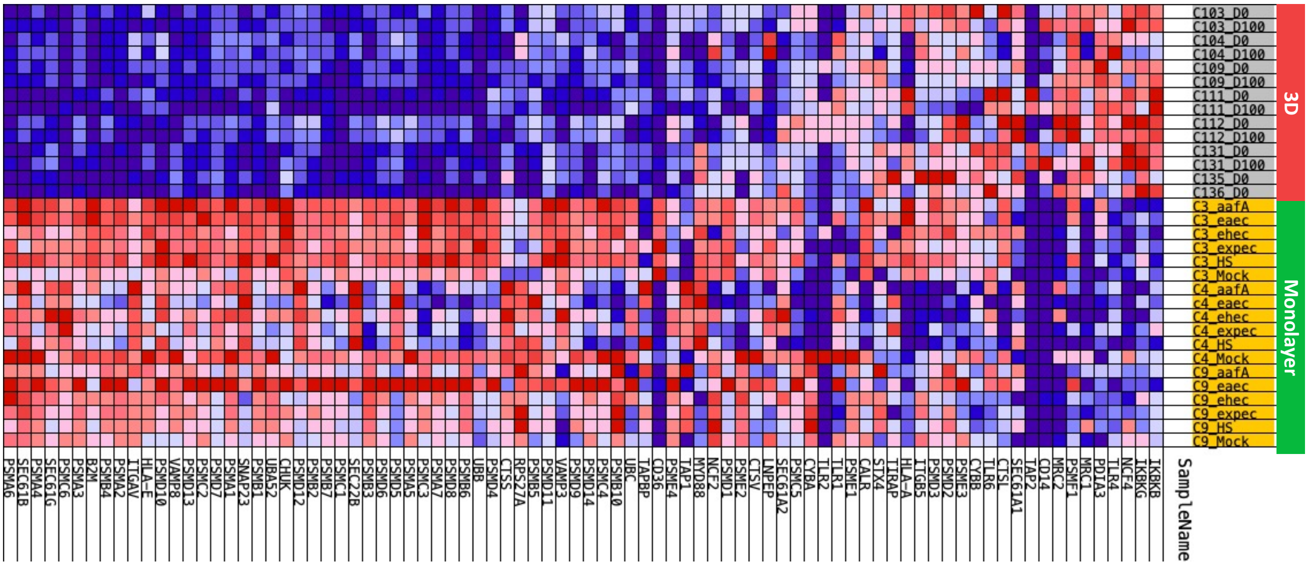

**Supplementary Figure 3. The antigen processing cross presentation gene set is upregulated in colonoid monolayers grown on Matrigel compared to differentiated 3D colonoids grown in Matrigel.** A.) PCA of the RNA-sequencing datasets for colonic enteroids: 3D-Matrigel (n=39), Monolayer-Matrigel (n=18) and Transwell-Collagen (n=12); B.) A dendrogram with agglomerative hierarchal clustering of the colonic gene set from RNA-sequencing. Branch length indicates degree of difference between samples; C.) GSEA showing an enrichment of the reactome antigen processing cross presentation gene set signature in colonoid monolayers grown on Matrigel when compared to 3D colonoids grown in Matrigel; D.) Heatmap of the reactome antigen processing cross presentation gene set showing an enrichment in colonoid monolayers grown on Matrigel (yellow) compared to 3D colonoids grown in Matrigel (gray);

**Supplementary Figure 4**

A.)

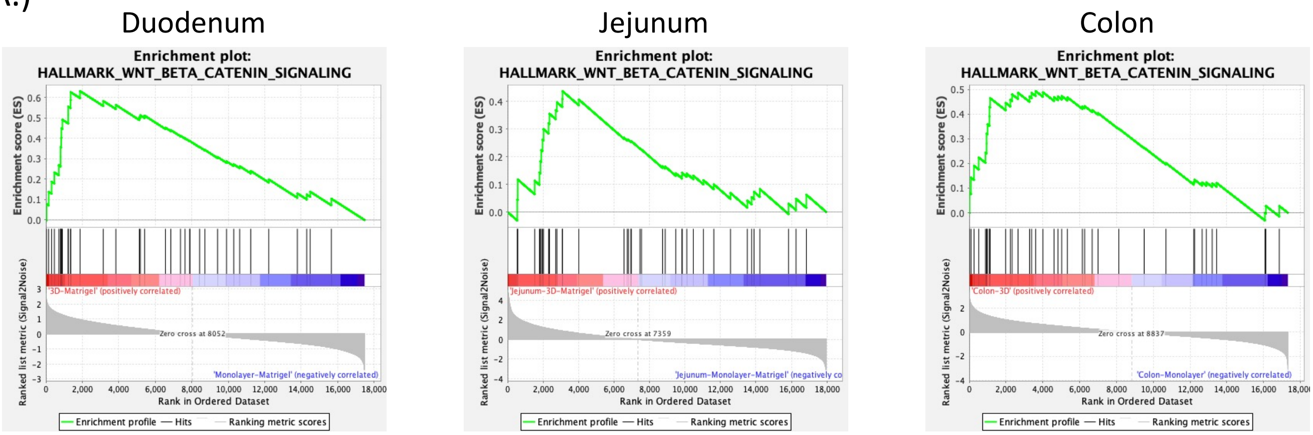


B.)

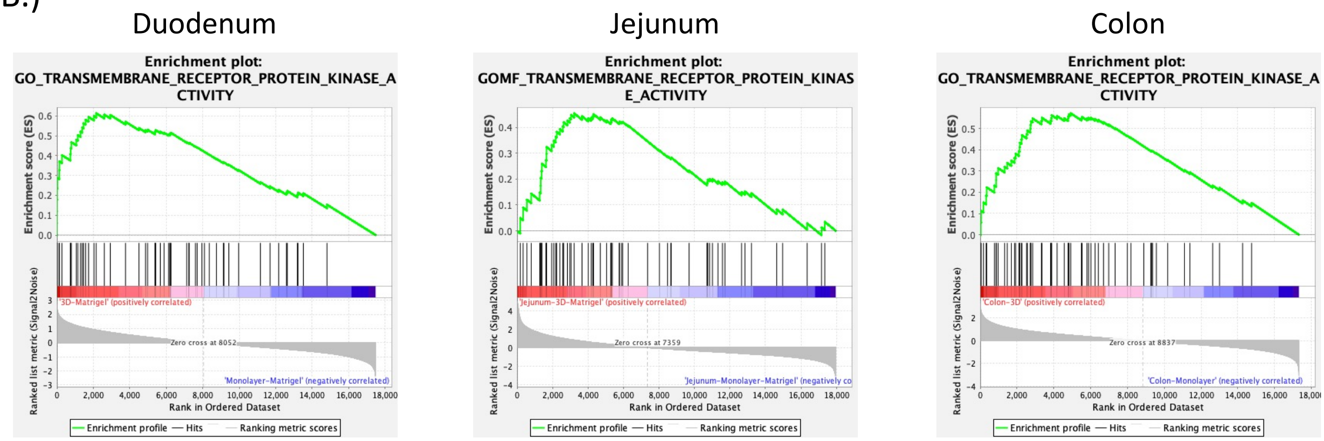


C.)

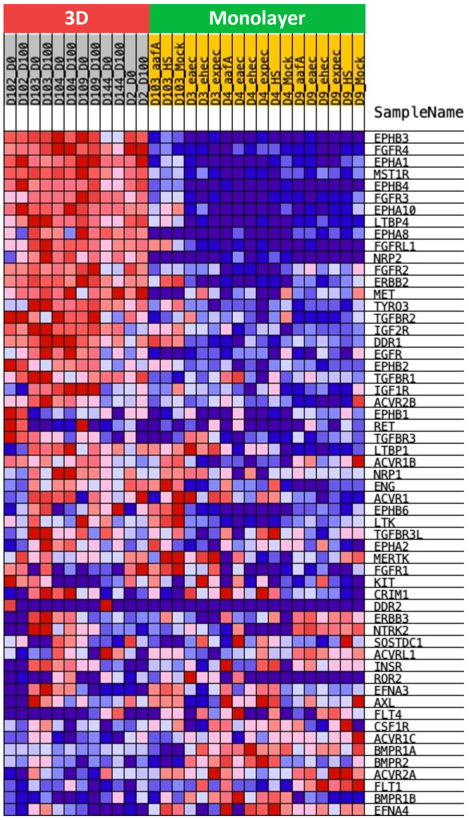

**Supplementary Figure 4. Wnt/β-catenin signaling and transmembrane receptor protein kinase activity genes are enriched in 3D intestinal organoids grown in Matrigel** compared to monolayers grown on Matrigel. A.) GSEA showing an enrichment of the hallmark wnt/β-catenin signaling get set signature in duodenal (left), jejunal (middle) and colonic (right) 3D organoids grown in Matrigel when compared to monolayers grown on Matrigel; B.) GSEA showing an enrichment of the GO transmembrane receptor protein kinase activity gene set signature in duodenal (left), jejunal (middle) and colonic (right) 3D organoids grown in Matrigel when compared to monolayers grown on Matrigel; C.) Heatmap of the transmembrane receptor protein kinase activity gene set showing an enrichment in 3D duodenal enteroids grown in Matrigel (gray) compared to Duodenal monolayers grown on Matrigel (yellow);

**Supplementary Figure 5**

A.) **Duodenum** B.) **Jejunum**

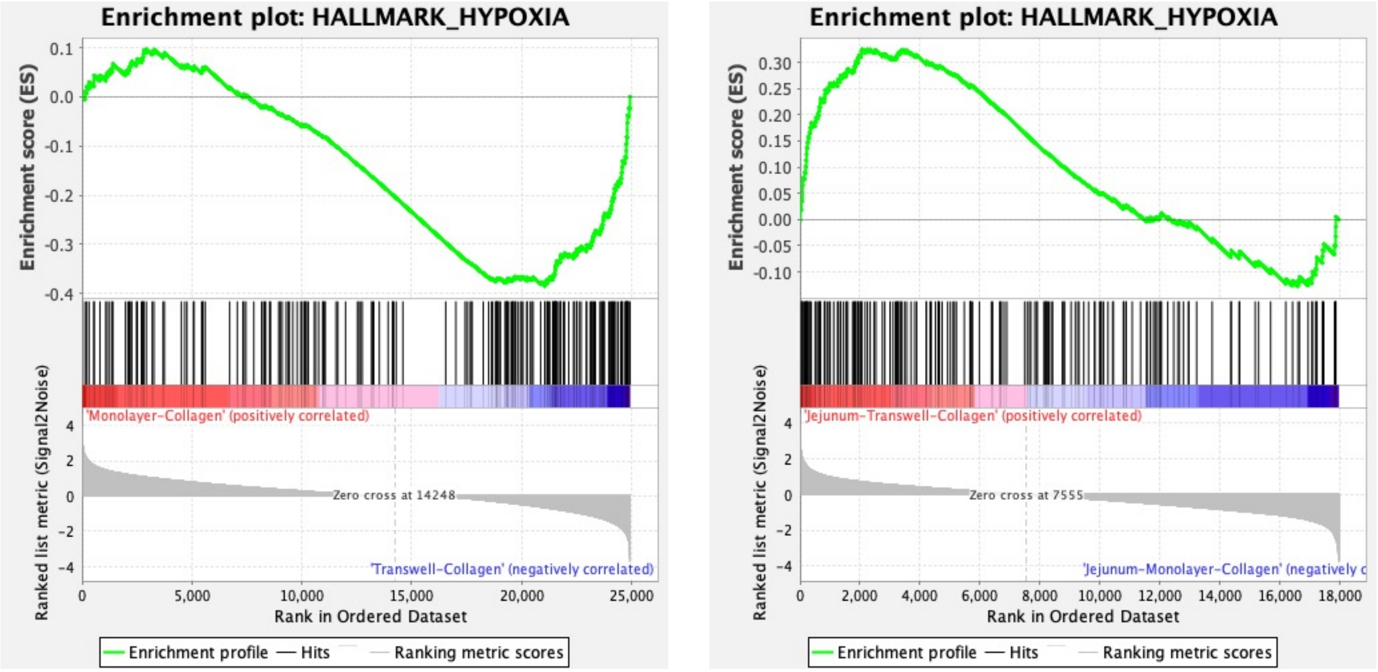


**Supplementary Figure 5. Hypoxia gene set is enriched in duodenal and jejunal transwells on Collagen compared to monolayers grown on collagen.** A & B.) GSEA showing an enrichment of the hallmark hypoxia get set signature in (A) duodenal (NES = -1.72 and FDR = 0.001) and (B) jejunal (NES = 1.78 and FDR = 0.003) transwells grown on collagen when compared to duodenal and jejunal monolayers grown on collagen;

**Supplementary Figure 6**

A.)

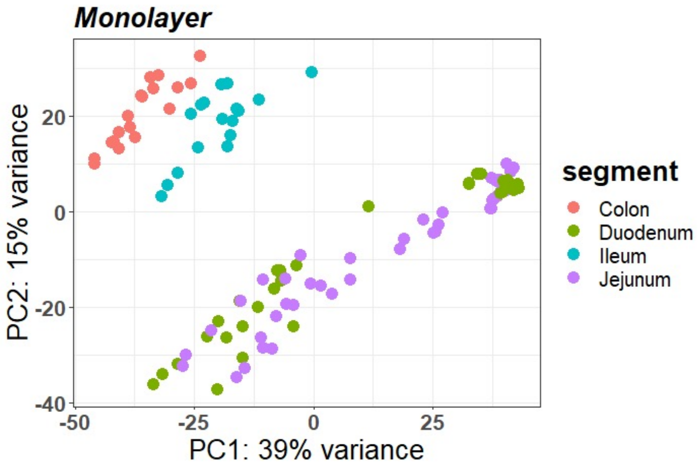


B.)

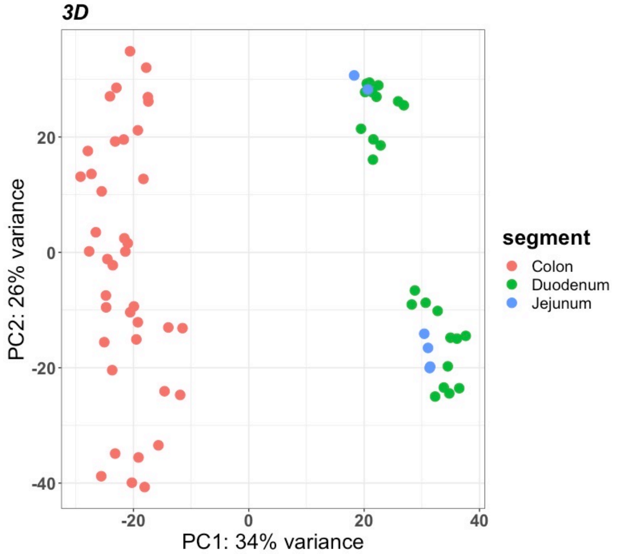


C.)

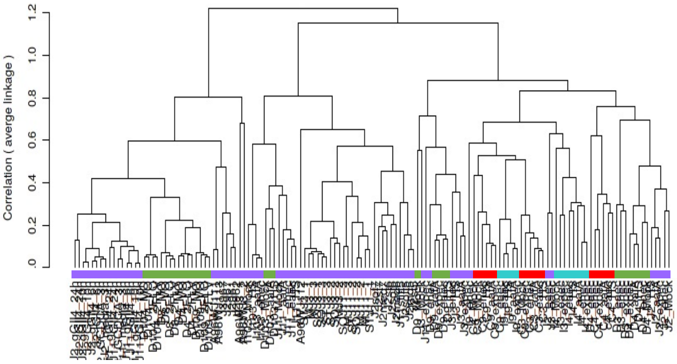


D.)

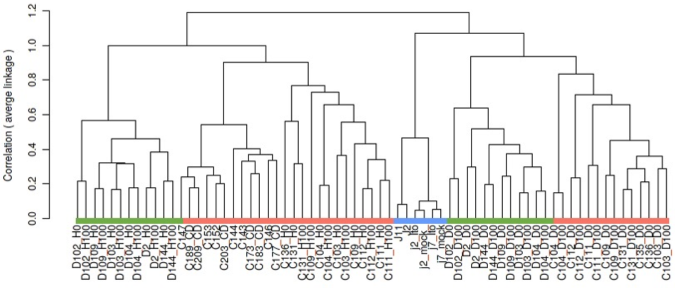


E.)

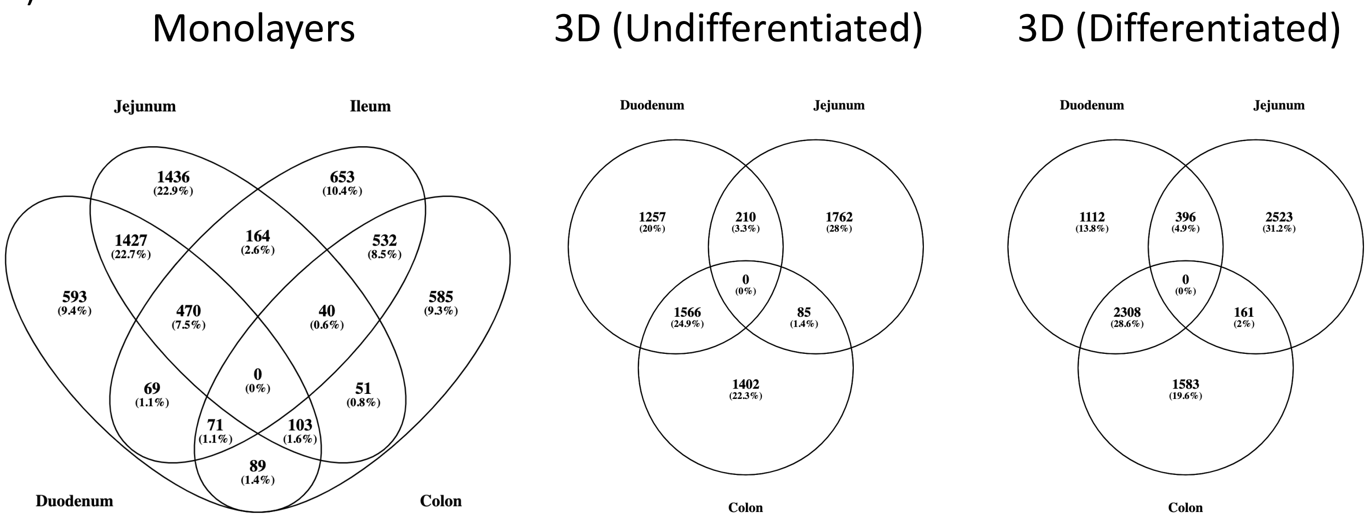

**Supplementary Figure 6. Transcriptionally segments vary in intestinal organoids grown in 3D and on monolayers.** A.) PCA of the RNA-sequencing datasets for enteroids on monolayers: Duodenum (n=26), Jejunum (n=69), Ileum (n=17), and Colon (n=18); B.) PCA of the RNA-sequencing datasets for enteroids on 3D: Duodenum (n=24), Jejunum (n=6), and Colon (n=39); C.) A dendrogram with agglomerative hierarchal clustering of the monolayer gene set from RNA-sequencing. Branch length indicates degree of difference between samples; D.) A dendrogram with agglomerative hierarchal clustering of the 3D gene set from RNA-sequencing. Branch length indicates degree of difference between samples; E.) Venn diagram displaying the overlap of differentially expressed genes (DEGs) between duodenal, jejunal, ileal and colonic intestinal organoids grown on Monolayers (Left) and in 3D (Undifferentiated (middle) and Differentiated (right)) . DEGs were were defined as any gene that was differentially enriched in the segment of interest compared to any other segment with DESeq2 (FDR ≤ 0.01 & Fold Change ≥2 and ≤ 0.5);

**Supplementary Figure 7**

A.)

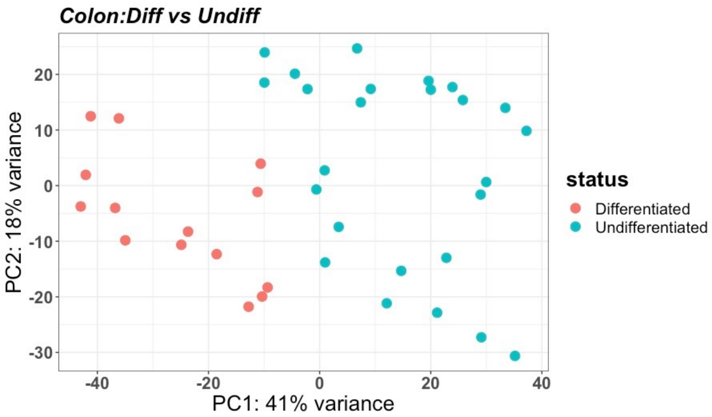


B.)

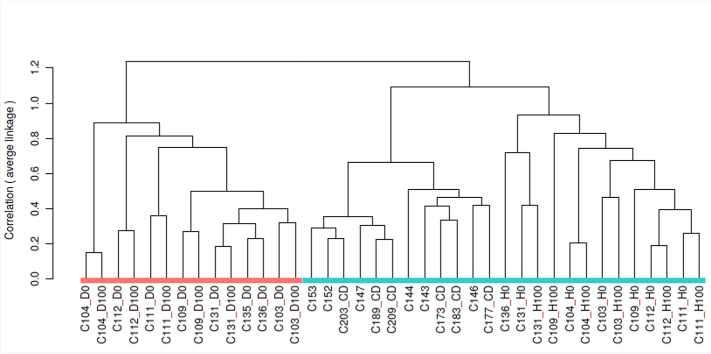


C.)

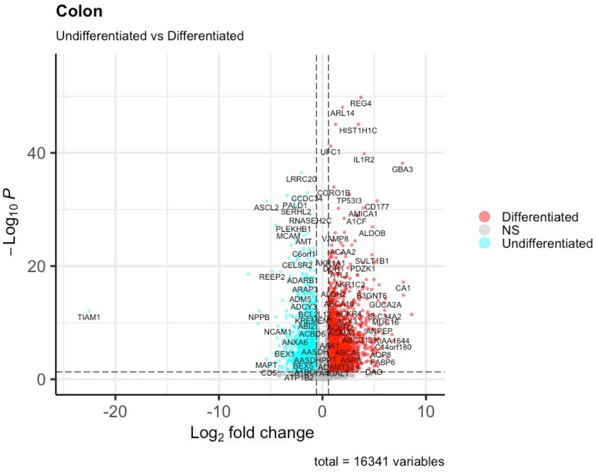


D.)

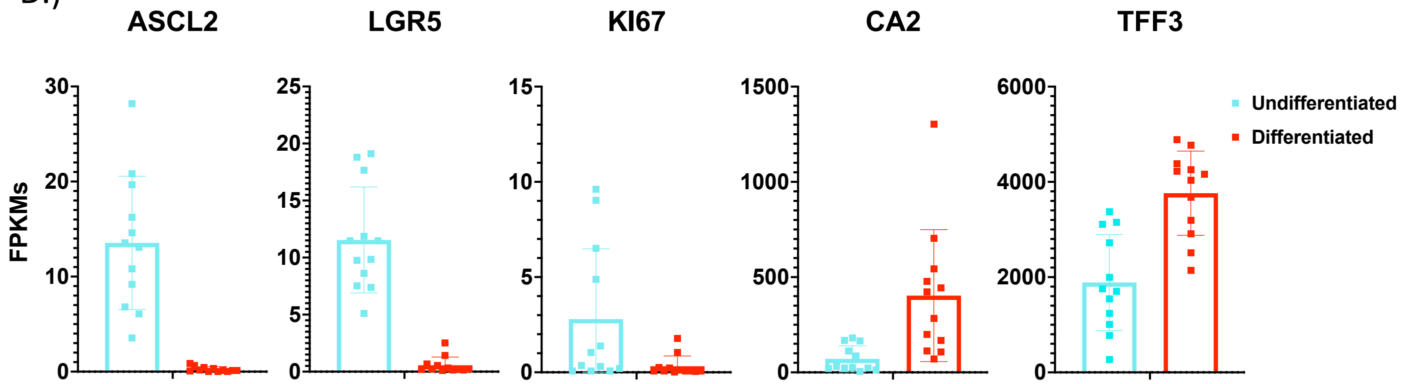


E.)

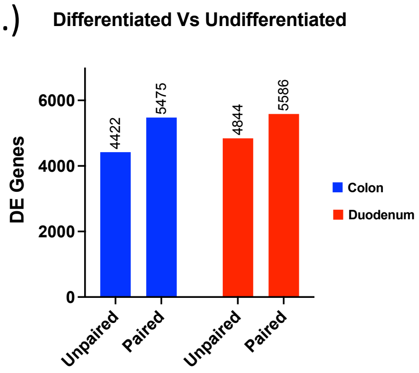

**Supplementary Figure 7. Differentiation media conditions drive changes in proliferation and differentiation markers in 3D colonoids.** A.) PCA of the RNA-sequencing datasets for 3D Colonic enteroids: Differentiated (n=14) and Undifferentiated (n=25); B.) A dendrogram with agglomerative hierarchal clustering of the 3D colonic gene set from RNA-sequencing. Branch length indicates degree of difference between samples; C.) Volcano plot of differentially expressed genes when comparing differentiated (red) and undifferentiated (cyan) 3D colonoids. Red/cyan dots indicate differentially expressed genes (FDR ≤ 0.01 and a foldchange ≥2 or ≤0.5) that are upregulated or downregulated (respectively) in differentiated (1338 genes) vs. undifferentiated (1418 genes) 3D colonoids; D.) Normalized FPKMs of undifferentiated (cyan) and differentiated (red) 3D colonoids for stem cell (*ASCL2*, *KI67* and *LGR5*) and differentiation (*CA2* and *TFF3*) markers; E.) The number of differentially expressed (DE) genes that were output from DESeq2 when comparing differentiated vs undifferentiated 3D duodenal enteroids (red) and colonoids (blue) as a result of using an unpaired (left) or paired (right) experimental design;

**Supplementary Figure 8**

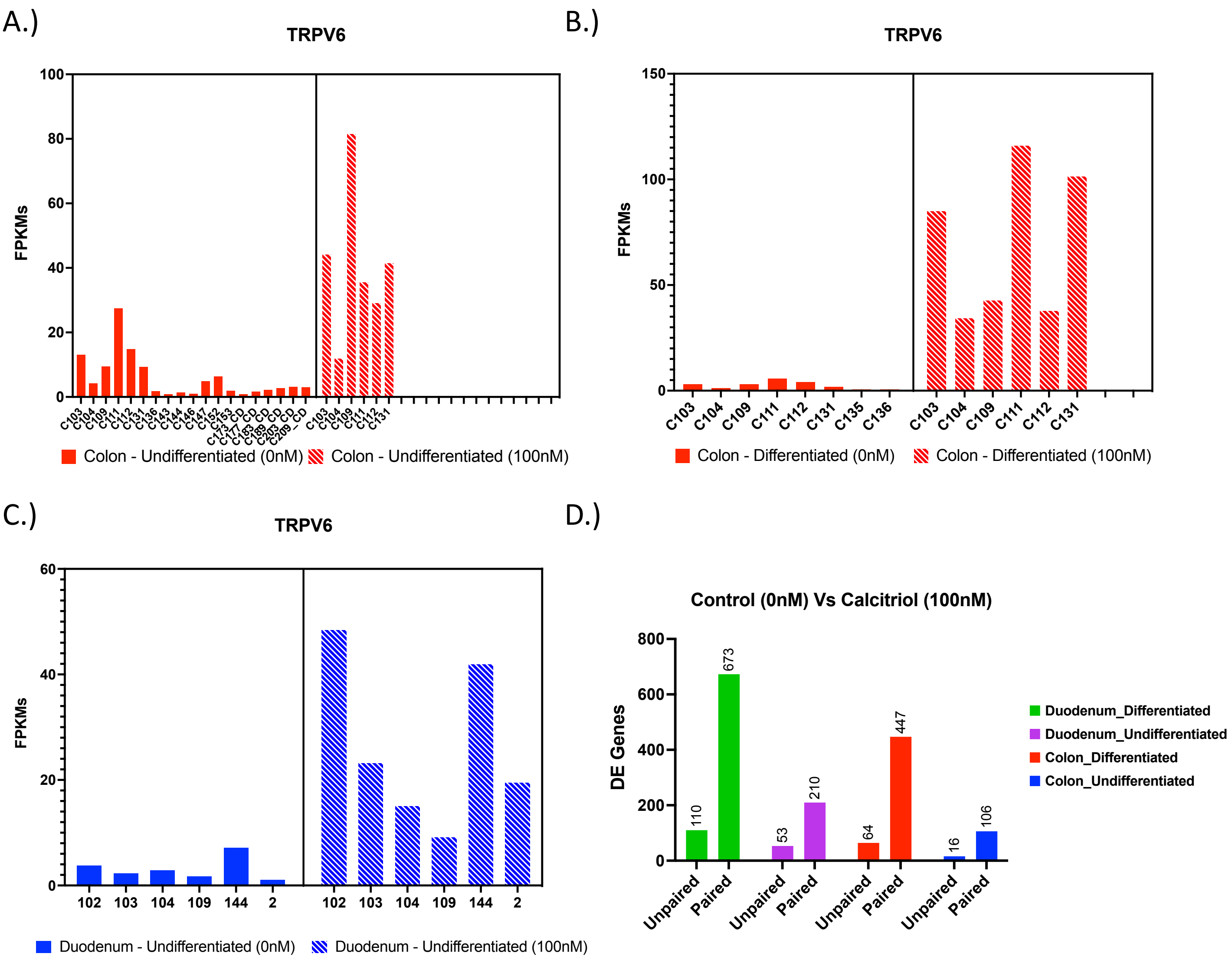


**Supplementary Figure 8. Patient-to-patient variability results in variable basal gene expression and response to stimuli of TRPV6 in differentiated and undifferentiated duodenal enteroids and colonoids.** A.) Normalized FPKMs of TRPV6 expression in undifferentiated patient derived colonoid lines comparing Control (solid) to 100nM Calcitriol treatment (dashed); B.) Normalized FPKMs of TRPV6 expression in differentiated patient derived colonoid lines comparing Control (solid) to 100nM Calcitriol treatment (dashed); C.) Normalized FPKMs of TRPV6 expression in undifferentiated patient derived duodenal enteroids lines comparing Control (solid) to 100nM Calcitriol treatment (dashed); D.) The number of differentially expressed (DE) genes that were output from DESeq2 as a result of using an unpaired (left) or paired (right) experimental design, for differentiated duodenal enteroids (green), undifferentiated duodenal enteroids (purple), differentiated colonoids (red) and undifferentiated colonoids (blue) treated with calcitriol (100nM) calcitriol compared to control (0nM);

**Supplementary Figure 9**

A.)

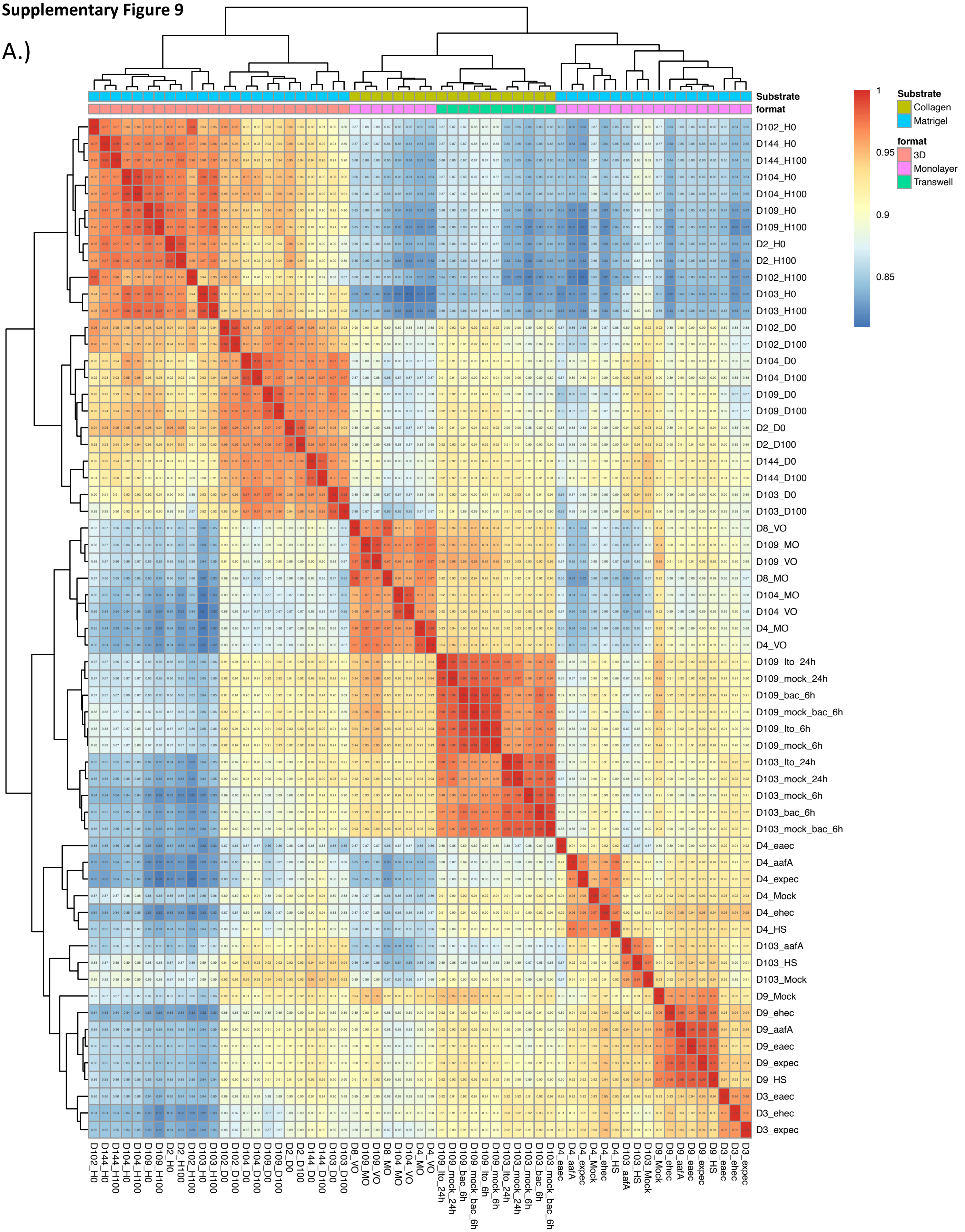


**Supplementary Figure 9. Correlation Matrix of Duodenal Enteroids.** A.) A Pearson correlation matrix of Duodenal enteroids

**Supplementary Figure 10**

A.)

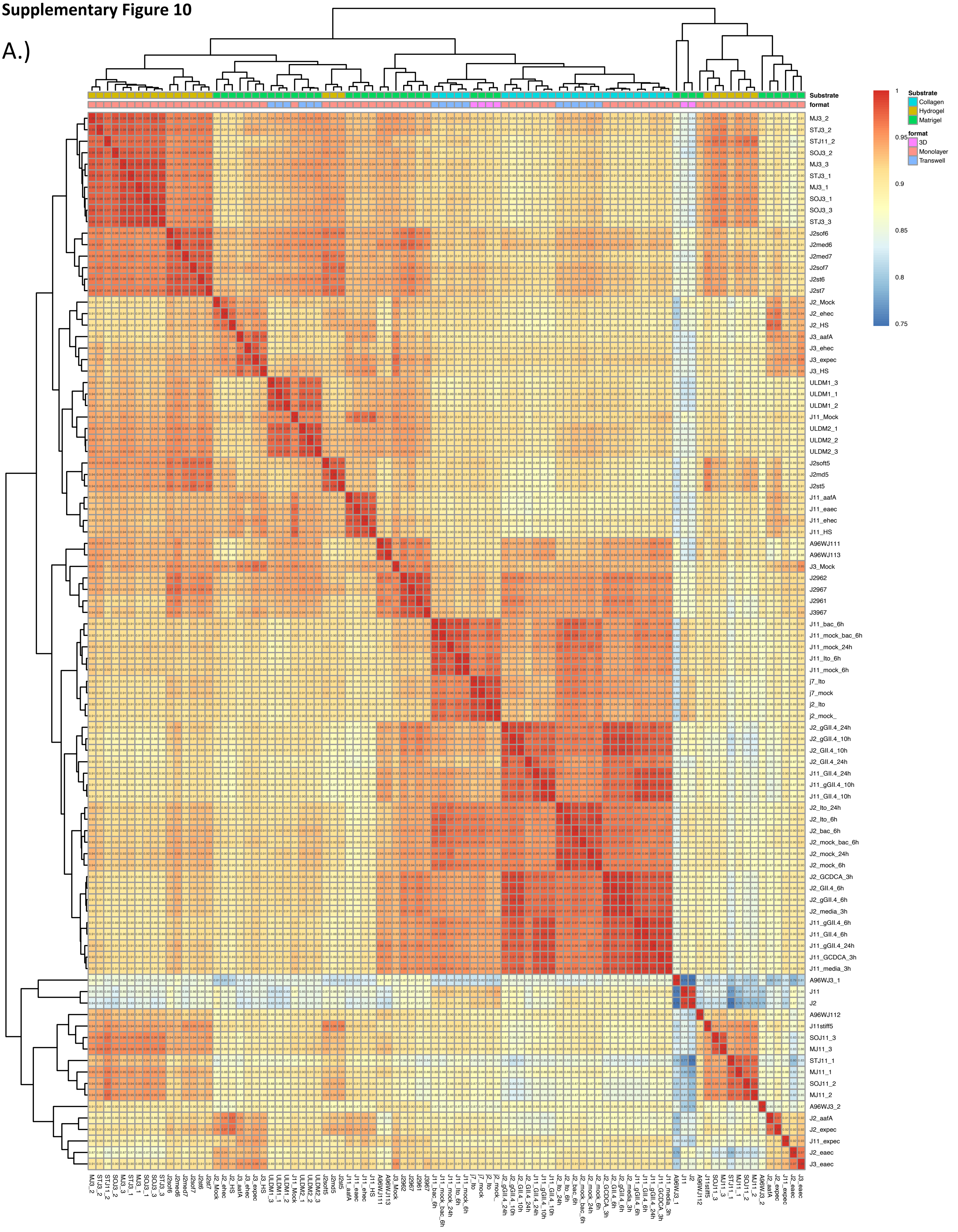


**Supplementary Figure 10. Correlation Matrix of Jejunal Enteroids.** A.) A Pearson correlation matrix of Jejunal enteroids

### Supplementary Figure 11

A.)

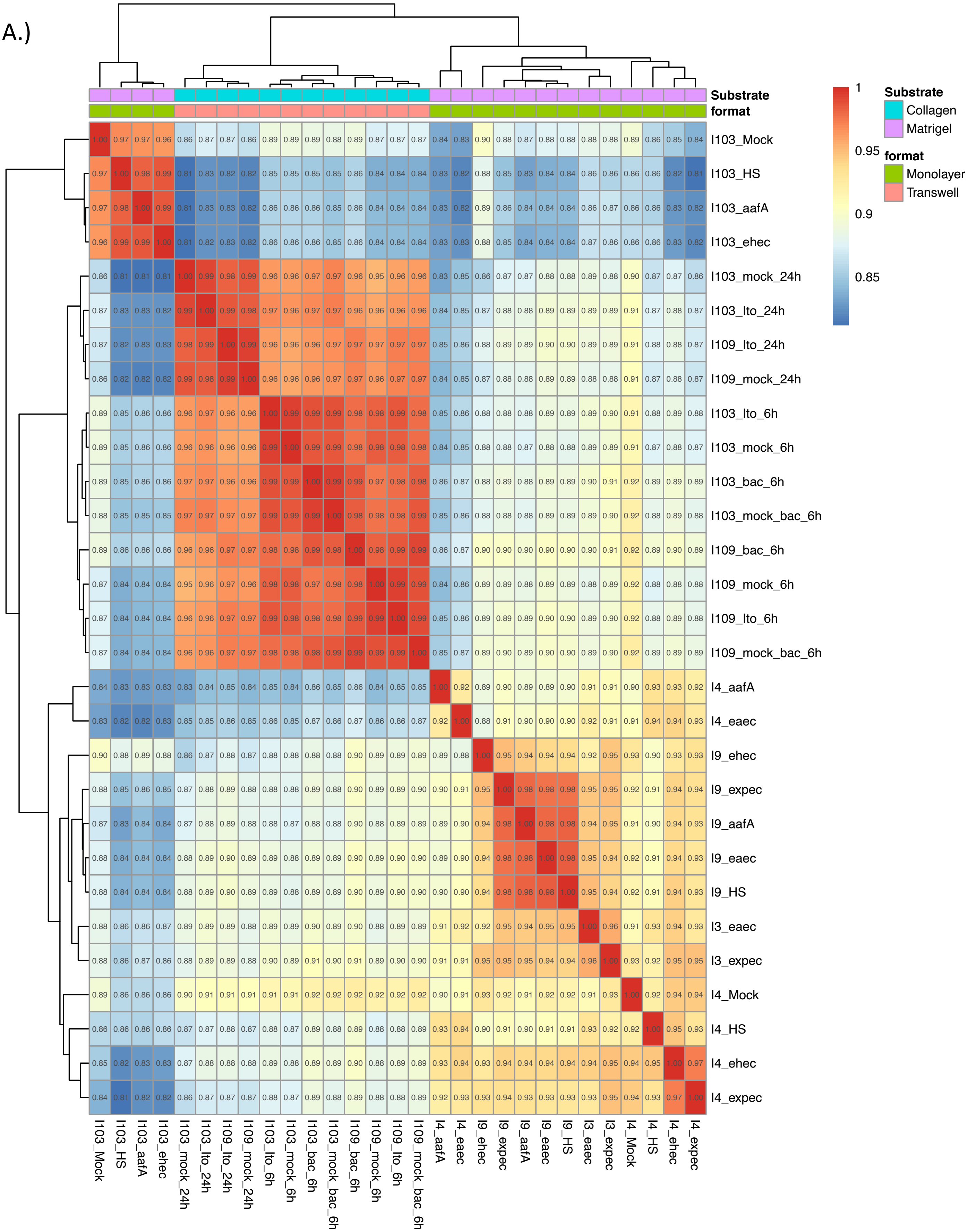


**Supplementary Figure 11. Correlation Matrix of Ileal Enteroids.** A.) A Pearson correlation matrix of Ileal enteroids.

**Supplementary Figure 12**

A.)

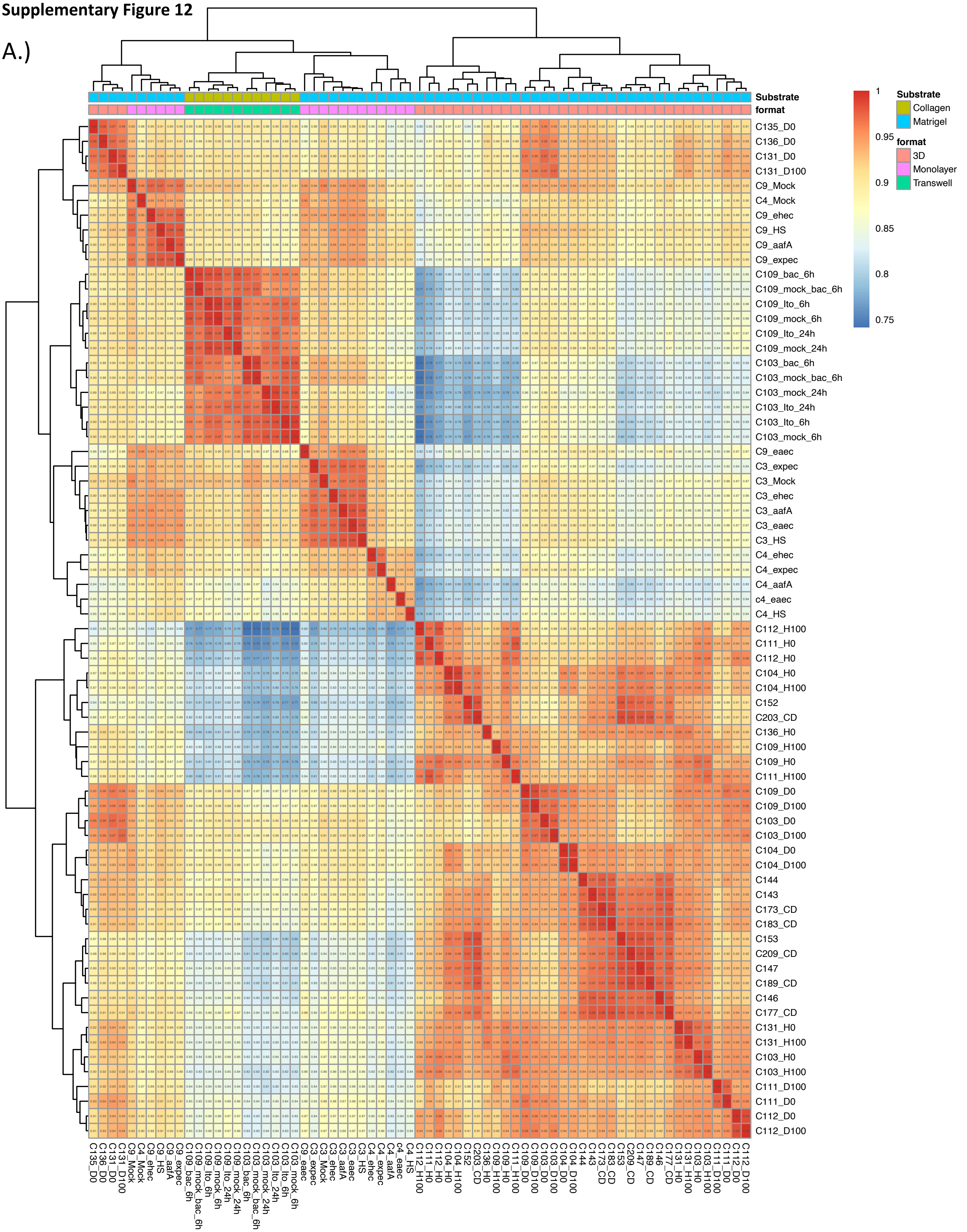


**Supplementary Figure 12. Correlation Matrix of Colonoids.** A.) A Pearson correlation matrix of Colonoids
